# Mandibular morphology clarifies phylogenetic relationships near the origin of crown birds

**DOI:** 10.1101/2025.09.26.678747

**Authors:** Abi H. Crane, Juan Benito, Albert Chen, Daniel T. Ksepka, Daniel J. Field

## Abstract

**Background:** The phylogenetic relationships of fossil birds near the origin of the avian crown group remain debated, in part due to a limited amount of character evidence from incomplete fossils. The avian lower jaw provides a potentially rich source of additional character data, yet fusion of the individual bony elements composing the avian post-dentary complex has impeded efforts to explore its phylogenetic signal. Here, we use high-resolution µCT-scanning to separate the individual bony elements of the mandibles of several immature crown birds and key fossil taxa, and use those data to assess support for alternative phylogenetic hypotheses for fossils near the origin of crown birds.

**Results:** We find that evidence from *Asteriornis* fails to support interpretations of derived mandibular similarities with palaeognaths, and instead strongly favours galloanseran, and specifically galliform, affinities. Our results also illustrate striking similarities in the architecture of the lower jaws between the toothed ornithurine *Ichthyornis*, Pelagornithidae and *Vegavis*, which, in addition to the absence of derived features linking them to Galloanserae, highlights questions regarding the phylogenetic position of these perennially controversial taxa.

**Conclusions:** Our data reveal new insight into patterns of morphological evolution near the origin of the avian crown group while raising new phylogenetic questions, emphasising the potential untapped value of detailed comparative investigations of morphological complexes such as the post-dentary complex of the mandible for informing the early evolutionary history of crown birds.

## Background

Neornithes (crown birds) are abundant and diverse in the modern world, with over 11,000 extant species (Gill & Donsker 2018). Whilst a large number of higher-level avian phylogenetic divergences appear to have taken place during a period of rapid diversification following the Cretaceous- Paleogene mass extinction, the earliest diverging neornithine lineages arose during the Cretaceous Period (Feduccia 2003; Clarke et al. 2005; Ericson et al. 2006; Prum et al. 2015; Ksepka et al. 2017; Field et al. 2020). There is a great deal of uncertainty surrounding the earliest evolution of Neornithes, with very little fossil evidence available from the Cretaceous with which to directly test inferences drawn from extant taxa. Given the paucity of the fossil record of early Neornithes (Mayr 2022), any known fossils from this time period prior to their extensive diversification are crucially important for informing our understanding of the ancestral condition of crown birds. Moreover, such fossils have potential to illuminate the extent to which crown birds diversified during the Cretaceous, and yield insight into the factors that enabled crown birds, uniquely among all dinosaurs, to survive the Cretaceous-Paleogene mass extinction. As such, developing a detailed understanding of the comparative morphology and phylogenetic position of available crown bird fossils from the earliest stages of the clade’s evolutionary history is critical.

The fossil record of Neornithes from the Mesozoic is extremely sparse. Over 50 genera of Mesozoic Euornithes are known, but almost all belong to stem-group birds, including the near-crown lineages Ichthyornithes and Hesperornithes (Bell & Chiappe 2020). Only two latest Cretaceous taxa are widely considered to be members of Neornithes: *Asteriornis maastrichtensis* and *Vegavis iaai*.

*Asteriornis*, the ‘wonderchicken’ from the latest Cretaceous of Belgium, is known from a nearly complete three-dimensionally preserved skull and some fragmentary post-cranial material. The phylogenetic position of *Asteriornis* is imperfectly resolved. Although all phylogenetic analyses to date have placed *Asteriornis* within Neornithes, its precise position therein remains uncertain. The combination of typically ‘anseriform’ and ‘galliform’ features in the holotype skull led Field et al. (2020) to suggest a probable phylogenetic position on the stem lineage of Galloanserae (the most exclusive clade uniting landfowl, Galliformes, and waterfowl, Anseriformes). Analyses from Field et al. (2020) variably placed *Asteriornis* as a stem-galloanseran, or stem-galliform depending on the phylogenetic method (maximum parsimony or Bayesian inference) applied. Parsimony analyses by Torres et al. (2021) instead recovered *Asteriornis* as a stem-palaeognath, albeit with weak support. The same position was recovered by Musser and Clarke (2024), though this was similarly weakly supported. The clade uniting *Asteriornis* and Palaeognathae was supported by a single synapomorphy in the Torres et al. (2021) dataset: a deeply caudally forked dentary with dorsal and ventral forks of approximately equal caudal extent. Using an updated version of the same dataset, Benito et al. (2022a) recovered *Asteriornis* variably as a stem-palaeognath (using parsimony) or crown anseriform (using Bayesian analyses). However, this dataset’s focus on stem birds may reduce its power to accurately resolve interrelationships among crown birds, given its limited sample of only four extant neornithines.

*Vegavis,* now thought to be slightly older than *Asteriornis* (Roberts et al. 2023) and thus the oldest known probable neornithine bird, is known from Vega Island, Antarctica (Clarke et al. 2005). It is known from multiple specimens which together preserve much of the skull, many post-cranial skeletal elements and a syrinx (Noriega & Tambussi 1995; Clarke et al. 2005; Clarke et al. 2016; West et al. 2019; Acosta Hospitaleche & Worthy 2021; Torres et al. 2025). It was initially described as a member of crown-group Anseriformes, deeply nested within Neornithes (Noriega & Tambussi 1995). However, the phylogenetic position of *Vegavis* has proven contentious, with alternative analyses resolving it in variable positions within Neornithes (Clarke et al. 2005; Clarke et al. 2016; Worthy et al. 2017; Field et al. 2020; Acosta Hospitaleche & Worthy 2021; Musser and Clarke 2024; Torres et al. 2025) or even outside of the neornithine crown group (O’Connor et al. 2011; McLachlan et al. 2017; Field et al. 2020).

Pelagornithidae is an enigmatic clade of Cenozoic fossil birds with an anatomy unique among Neornithes. The group persisted through most of the Cenozoic era, with the earliest known pelagornithid known from the early Palaeocene (61.5-62 Ma; Mayr et al. 2021) and the latest from the late Pliocene (∼2.5 Ma: Boessenecker & Smith 2011). In addition to prominent bony pseudoteeth lining the upper and lower jaws, later representatives of this group are notable for their large size; at 6.4m, the late Oligocene *Pelagornis sandersi* had the largest wingspan of any known volant bird (Ksepka 2014). Despite being well-represented in the fossil record, the phylogenetic position of Pelagornithidae has been controversial. As with both *Vegavis* and *Asteriornis*, pelagornithids have been considered putative total-clade galloanserans, with some analyses resolving them as sister to Anseriformes (Bourdon 2005; Bourdon 2011; Field et al. 2020; Musser and Clarke 2024), sister to Galloanserae (Mayr 2011), or sister to Neognathae (Houde et al. 2023). Other analyses have resulted in pelagornithids being placed in a polytomy with Galloanserae and Neoaves (Mayr et al. 2021).

The avian mandible, especially the anatomy of its individual constituent bones, has been critically understudied, to the extent that even detailed morphological studies of extant birds generally do not consider the individual bones of the mandible (e.g., Jones et al. 2019). This dearth of knowledge is primarily a result of the high degree of fusion of individual osteological components in the mandibles of adult birds (Jollie 1957; Plateau & Foth 2020) which can render them completely indistinguishable (Hogg 1983). Mandibular fusion occurs largely post-hatching, with sutural obliteration progressively obscuring the identity of individual bones (Jollie 1957; Bellairs et al. 1960). In extant birds, the bones comprising the ‘post-dentary complex’ (the angular, surangular, prearticular and articular) are among the first mandibular bones to fuse (Hogg 1983; Plateau & Foth 2020). However, the caudal boundary of the dentary is often possible to discern even in adult birds (Pycraft 1900); as such, in fossil birds, it is often possible to distinguish between the dentary and the post- dentary complex, but not among the individual components of the post-dentary complex (e.g., Chiappe et al. 2007; Wang et al. 2021).

Despite this lack of insight into the comparative morphology of the mandible among extant birds, multiple aspects of avian mandibular morphology, such as the presence and morphology of retroarticular processes, are recognised to be highly phylogenetically informative. An enlarged retroarticular process represents one of very few osteological synapomorphies diagnosing crown Galloanserae (Mayr 2017; Field et al. 2020), uniting the otherwise osteologically divergent Galliformes and Anseriformes (Olson & Feduccia 1980). Given that the detailed mandibular morphology of extant birds remains relatively unknown, other phylogenetically important features of the mandible may await recognition.

In recent years, the first well-preserved mandibles of Cretaceous probable neornithines have come to light, alongside new pelagornithid specimens which offer further insight into their unusual mandibular morphology (Mayr & Rubilar-Rogers 2010; Mayr et al. 2021). To date, however, the detailed comparative morphology of their mandibles has provided limited insight into their phylogenetic affinities. Here we reinvestigate the mandibular morphology of *Asteriornis, Vegavis*, *Ichthyornis* and the early pelagornithid *Dasornis toliapica* (formerly ‘*Odontopteryx toliapicus*’ (Owen, 1873; Bourdon et al. 2010). We make comparisons with the mandibular anatomy of major crown bird lineages, characterized for the first time using high-resolution µCT scans of juvenile specimens.

## Methods

### Taxon and specimen selection

The mandibular anatomy of the following phylogenetically controversial fossil specimens was examined in detail: *Asteriornis* (NHMM 2013 008), *Vegavis* (AMNH FARB 30899) and *Dasornis* (NHMUK 44096).

Juvenile representatives of neornithines were examined to characterise the morphology of the following groups: Palaeognathae (*Struthio camelus, Dromaius novaehollandiae, Tinamus solitarius*), Anseriformes (*Chauna chavaria, Thalassornis leuconotus, Anser fabalis*), Galliformes (*Megapodius pritchardii, Gallus gallus*), and representatives of Neoaves bearing intraramal joints (*Morus bassanus, Diomedea exulans*). Skeletally immature specimens were selected for investigation since the bones of the avian mandible become more fused during post-hatching ontogeny (Jollie 1957; Webb 1957; Hogg 1983). Some features of the avian mandible are not present or differ morphologically in juvenile specimens when compared with adults (e.g., the galloanseran retroarticular process), so comparisons with unsegmented skeletally mature individuals were also made, where appropriate. For anatomical comparisons with the crownward stem-bird, *Ichthyornis*, we used several mandible specimens of varying levels of completeness and fusion of the post-dentary complex: YPM 1450, BHI 6421, KUVP 119673 and AMNH FARB 32773 (Field et al. 2018; Torres et al. 2021).

### CT scanning and segmentation

µCT scanning of extant taxa and *Asteriornis* was performed at the Cambridge Biotomography Centre (CBC). Scanning of *Dasornis toliapica* (NHMUK 44096) was performed at the Natural History Museum, London. Scans of *Ichthyornis* were originally performed at the University of Texas High-Resolution CT Facility (UTCT) (KUVP 119673 and BHI 6421) and the Center for Nanoscale Systems at Harvard (YPM 1450). A digital mesh of the predentary of *Ichthyornis* (AMNH FARB 32773) was obtained from the supplementary material of Torres et al. (2021), and CT data and digital meshes of *Vegavis* (AMNH FARB 30899) were obtained from Torres et al. (2025).

All scanned specimens were digitally segmented and rendered in VGSTUDIO MAX 3.4 (Volume Graphics, Heidelberg, Germany). 3D surface meshes were created in VGSTUDIO MAX and exported into Autodesk Maya 2023, where digital 3D reconstruction was performed, see below.

### Anatomical description

We use osteological nomenclature following Baumel and Witmer (1993), though we choose to use the term ‘surangular’ instead of ‘supra-angular’ as its use has become well-established in recent decades (e.g., Elzanowski 1999; Clarke 2004; Field et al. 2018).

Following Baumel and Witmer (1993), we use angulus mandibulae (‘mandibular apex’) to describe the dorsal prominence of the post-dentary complex formed by the surangular. This structure corresponds to the coronoid process and serves as the attachment point for the external adductor muscles of the mandible in most neornithine birds. In contrast, the coronoid process of Anatidae and megapode galliforms forms as a laterally projecting outgrowth of the surangular (Schneider 2024). The mandibular apex is often referred to as the coronoid process in descriptions of fossil taxa (including those of fossil anseriforms; e.g., Tambussi et al. 2019; Houde et al. 2023; Musser & Clarke 2024). We refer to this position as the mandibular apex to avoid confusion when making anatomical comparisons between birds in which this structure is synonymous with the coronoid process and those where the coronoid process is a separate structure.

In addition to the taxa selected for CT scanning, further anatomical comparisons to extant taxa were made with reference to specimens held in the University of Cambridge Museum of Zoology (UMZC). Additional anatomical comparisons were made with reference to surface scans of an undescribed pelagornithid from the Miocene of Peru (MUSM 1677).

### 3D reconstruction

The articulated mandible of *Asteriornis* is shattered into many adjacent fragments of varying sizes; every fragment that was large enough to segment and confidently identify as part of the mandible was included as part of our 3D reconstruction. Each fragment that could be separated along lines of breakage was exported as a separate mesh to enable maximum flexibility in the reconstruction (see Supplementary Fig. 1 for NHMM 2013 008 segmentation). The digital meshes were exported to Autodesk Maya 2023, where they were manipulated and re-oriented to reconstruct their original anatomical position. This was achieved by matching broken edges to create cohesive bones where possible and referencing more complete parts of each mandibular ramus to maintain symmetry.

A complete model of the mandible of *Ichthyornis* was constructed using digital meshes from four specimens (YPM 1450, BHI 6421, KUVP 119673, AMNH FARB 32773; Supplementary Fig. 2) in Autodesk Maya 2023. The specimens of *Ichthyornis* represent individuals of different sizes, so meshes were scaled relative to BHI 6421 to create a single cohesive anatomical model.

A complete model of the mandible of *Vegavis* was constructed using two digital meshes from AMNH FARB 30899 (Torres et al. 2025): the well-preserved caudal part of the left mandibular ramus and a mirrored mesh of the rostral portion of the right mandibular ramus.

## Results

### Asteriornis

#### General morphology

The mandibular symphysis of *Asteriornis* is unusual and is not directly comparable to that of any extant neornithines examined. The symphysis is deep both dorsoventrally and rostrocaudally, forming a distinctive ‘scooped’ morphology (Fig. 1). The symphysis does not resemble the upturned occlusal surface typical of palaeognaths (Fig. 2B; Nesbitt & Clarke 2016), the ventrally slanted and pointed tip of galliforms (Fig. 2E), the round but ventrally pointed rostral end of anhimids, or the flat and broad spatulate symphysis seen in Anatidae (Fig. 2D). This unusual morphology of *Asteriornis* has not previously been recognised, given that the left side of the symphysis is fractured and thus the true shape of the symphysis in the fossil itself is obscured (Field et al. 2020; Torres et al. 2021).

**Figure 1.**
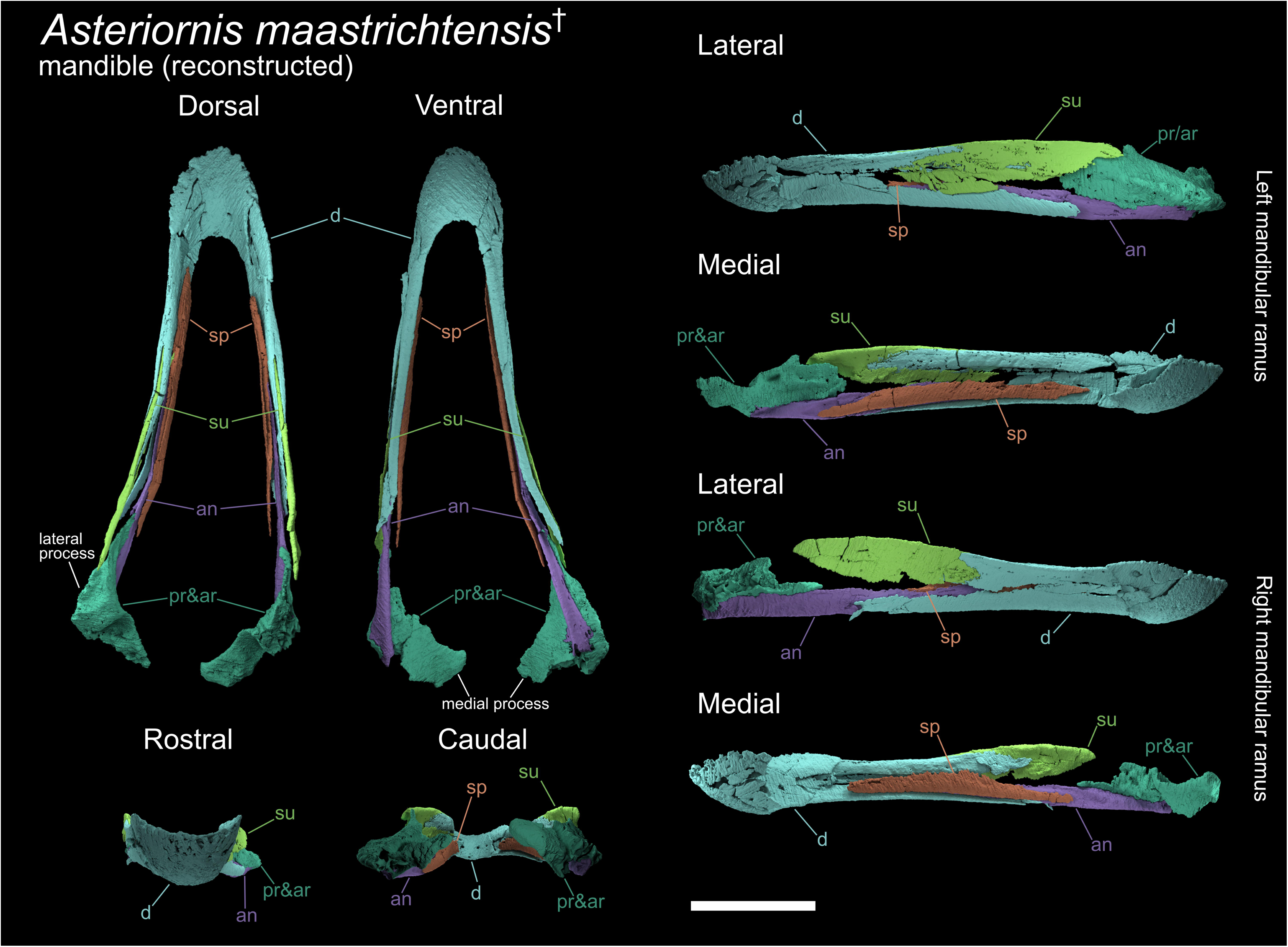
Digitally reconstructed mandible of *Asteriornis*. Whole mandible shown in dorsal, ventral, rostral and caudal views. Left and right rami shown separately in medial and lateral views. Prearticular and articular could not be distinguished, displayed as prearticular/articular combined. Abbreviations: an, angular; ar, articular; d, dentary; pr, prearticular; sp, splenial; su, surangular. Scale bar equals 10mm.

**Figure 2.**
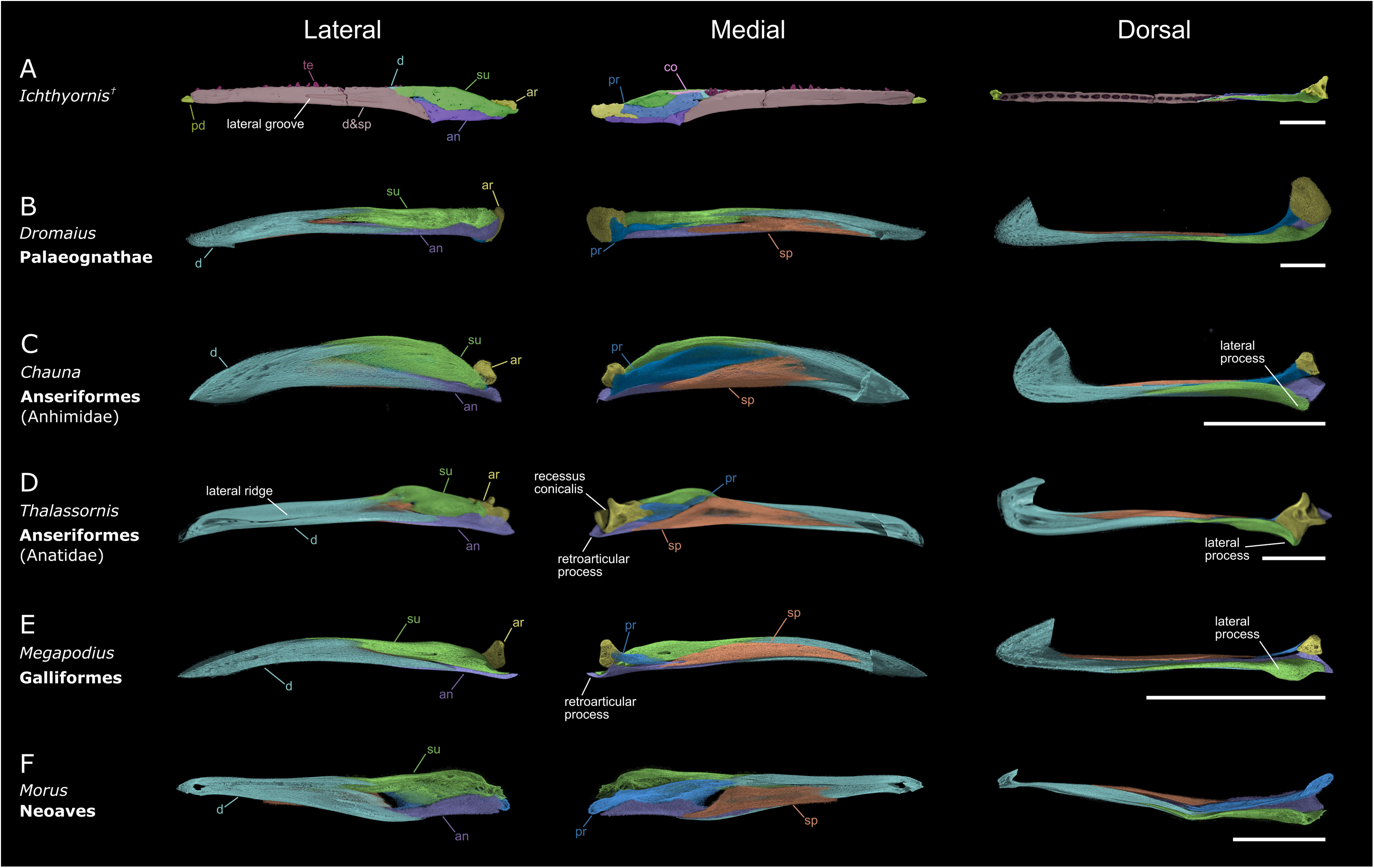
Mandibles of *Ichthyornis* and representative immature crown-group birds. Left mandibular rami shown in lateral, medial and dorsal views. A) Digitally reconstructed mandible of *Ichthyornis* created from digital meshes of specimens BHI 6421, KUVP 119673, YPM 1450 and AMNH FARB 32773 (for details on fossil segmentation, see Supplementary Fig. 2). The dentary and splenial bones of Ichthyornis are highly fused and impossible to differentiate for most of their length (see Supplementary Fig. 7); as such, the dentary/splenial are not distinguished in this figure and coloured accordingly. Additional abbreviations: co, coronoid; pd, predentary; te, teeth. B) *Dromaius novaehollandiae* (Palaeognathae). C) *Chauna chavaria.* D) *Thalassornis leuconotus* (Anseriformes). E) *Megapodius pritchardii* (Galliformes). F) *Morus bassanus* (Neoaves); this specimen is at an early ontogenetic stage and does not exhibit an ossified articular bone. Further immature neornithine mandibles are figured in the supplementary material (Supplementary Figs. 3, 4, 5, 6). Scale bar equals 10mm.

The mandibular rami of *Asteriornis* are extremely narrow mediolaterally relative to their dorsoventral depth. There is a slight increase in dorsoventral depth towards the caudal end of the mandible, peaking near its caudal terminus. This resembles the typical morphology of galliforms and Anhimidae (Fig. 2C, E; Mayr et al. 2023), in which the mandibular ramus maintains a nearly equal dorsoventral depth along its entire length, and contrasts with the shape of the more derived anatid mandible (Fig. 2D), which deepens caudally with a prominent mandibular apex.

The dorsal and ventral margins of the *Asteriornis* mandible are approximately flat and parallel to one another along their length, with little discernible curvature (Fig. 1 lateral, medial). This contrasts with the galliform and anhimid condition, which is curved with a convex dorsal and concave ventral margin (Fig. 2C,E). Although not as pronounced, the palaeognath mandible is similarly curved (Fig. 2B). In this respect, the mandible of *Asteriornis* is similar to that of anatid anseriforms among galloanserans, though this generalized mandible shape is widespread in birds (Fig. 2D).

#### Dentary

*Asteriornis* does not exhibit the rostrocaudally-oriented lateral ridge of the dentary seen in Anatidae (Fig. 2D), which appears to be apomorphic for this group.

The dentary of *Asteriornis* exhibits a forked caudal end. This morphology has been previously interpreted as similar to the deeply forked condition of palaeognaths, with dorsal and caudal ends of the fork of approximately equal caudal extent (Torres et al. 2021). However, reconstruction of the mandible of the right mandibular ramus of *Asteriornis* illustrates that the caudal fork of the dentary is substantially shallower than would be interpreted from the more fragmentary left side (Supplementary Fig. 1, left lateral). In the reconstructed mandible, the caudal forking of the right dentary is relatively shallow (Fig. 1 right lateral), with the dorsal prong dorsoventrally shallower (extending less far caudally) than the ventral prong. This morphology closely resembles the caudal end of the dentary in galliforms and anhimids (Fig. 2C,E), in which the degree of forking is similarly asymmetric, with a substantially smaller dorsal process, and contrasts with the palaeognath condition, which exhibits prongs that are much more similar in length (Fig. 2B, Supplementary Fig. 3), a condition also present in certain Neoaves (e.g., *Morus*, Supplementary Fig. 4).

A rostrally projecting process of the surangular runs along the midline of the dorsal edge of the mandible in *Asteriornis*, dividing the mandible mediolaterally. This mediolateral splitting of the dentary by the surangular occurs well rostral to the abovementioned forked caudal end of the *Asteriornis* mandible, resembling the condition in galliforms, whereas in palaeognaths and the surveyed neoavians the surangular divides the mandible mediolaterally at approximately the point of divergence of the dorsal and ventral processes of the forked caudal mandible. Despite a comparable rostral extent of the surangular in palaeognaths and *Asteriornis*, these observations demonstrate that the dentary of *Asteriornis* is substantially more similar to that of galliforms and anhimids than it is to palaeognaths.

#### Splenial

The left splenial of *Asteriornis* (Fig. 1 left medial) is preserved *in situ*, and the right splenial is disassociated from the rest of the mandibular ramus. The overall shape of the splenial is rod-like and has an approximately consistent dorsoventral depth for most of its length (Fig. 1 left medial). This shape is similar to that in galliforms and palaeognaths and is very unlike the distinctive, large, triangular splenial of extant anseriforms (Fig. 2C,D). As noted by Field et al. (2020), the splenial of *Asteriornis* is unfused to the rest of the mandible along its entire length. This condition is observed in adult representatives of Palaeognathae (Houde & Haubold 1987; Nesbitt & Clarke 2016) and the galliform subclade Megapodiidae (Field et al. 2020). In most other extant birds, the splenial is fused to the dentary along at least the rostral portion of its length in adult birds.

The splenial extends less far rostrally than in palaeognaths (Fig. 2B), in which it reaches the mandibular symphysis in some taxa (e.g., *Tinamus*; Supplementary Fig. 3C) (Houde & Haubold 1987; Nesbitt & Clarke 2016). The relatively caudal position of the splenial in *Asteriornis* falls within the range of variation encompassed by extant galloanserans (Fig. 2C-E), not reaching the mandibular symphysis (as is also the case in most other non-palaeognath birds). . The splenial does not contribute to the ventral margin of the mandibular ramus, as occurs in Anseriformes, some palaeognaths and *Ichthyornis* (Fig. 2A-D).

The splenial of *Asteriornis* does not appear to contact the prearticular; even when poor preservation is considered, their relative positions illustrate that contact between these elements is unlikely. This condition is exhibited by galliforms and some palaeognaths (e.g., *Dromaius*; Fig. 2B,E), whereas anseriforms exhibit an apparently derived condition with extensive contact between the splenial and prearticular (Fig. 2C,D).

#### Post-dentary complex

The dorsal margin of the *Asteriornis* surangular is primarily straight, running parallel to the rest of the mandibular ramus, with only a slight degree of curvature (Fig. 1) and an indistinct mandibular apex. This is similar to the condition in galliforms, palaeognaths and anhimids (Fig. 2B,C,E) and contrasts with the prominent mandibular apex of the anatid surangular (Fig. 2D). Notably, the surangular of *Asteriornis* appears to be slightly dorsally directed, and bounded ventrally by a thick and robust articular/prearticular. Thus, the surangular and the angular are widely separated from each other in lateral view and do not appear to contact caudally. Although likely influenced by taphonomic deformation, if genuine, this feature would distinguish *Asteriornis* from all examined crown and stem- birds (Figs. 1 & 2). Although this condition is apparent in both mandibular rami (Fig. 1; Supplementary Fig. 1), the poor preservation of the caudal end of both lower jaws precludes a fully confident assessment of this arrangement. Considering the limited degree of fusion of the *Asteriornis* lower jaw, it is not inconceivable that the post-dentary elements were slightly taphonomically shifted, with the incomplete preservation of the articular/prearticular preventing a more accurate reconstruction of how these elements were arranged in life.

The lateral mandibular process of *Asteriornis* forms a low but robust and elongate ridge, most closely resembling the galliform condition (Crane et al. 2025). By contrast, the lateral mandibular process of palaeognaths is only slightly developed (e.g., in *Struthio*) or non-existent (e.g., in *Dromaius* and *Tinamus*: Fig. 2B, Supplementary Fig. 3) and that of anseriforms is rostrocaudally short and laterally prominent (Fig. 2C,D).

The caudal ends of the mandible are too poorly preserved to confirm or reject the possible presence of a galloanseran-like retroarticular process (Crane et al. 2025). The angular extends all the way to the caudalmost preserved point of the mandible (and thus could have contributed to the formation of the retroarticular process if one were present), whereas we interpret the surangular as terminating rostral to the position of the hypothetical retroarticular process. Thus, if present, the retroarticular process would have been composed of only the angular as in anseriforms (Fig. 2C,D), rather than of a combination of both surangular and angular as in galliforms (Fig. 2E).

As preserved, the rostral portion of the *Asteriornis* prearticular is most comparable to that of galliforms and palaeognaths, though it exhibits a slightly greater degree of overlap with the splenial than in either of these extant groups. Given that the rostral portion of the prearticular/articular is poorly preserved, it is likely that the true rostral extent of the prearticular would be greater and thus intermediate between that of anseriforms and galliforms/palaeognaths. The shape of this rostral portion is deep, comparable to the robust rostral termination of the prearticular of anseriforms (Fig. 2C,D medial) rather than the narrower, pointed termination observed in palaeognaths and galliforms (Fig. 2B medial; Fig. 2E medial). *Ichthyornis* exhibits a similarly dorsoventrally deep prearticular (Fig. 2A medial) which is rostrally extensive, overlying the dentary/splenial region much like the condition in anseriforms.

There is no evidence of a prominent recessus conicalis in *Asteriornis* (Fig. 1 caudal). In Anatidae, the recessus conicalis takes the form of a deep and conical hollow on the caudal surface of the medial mandibular process (Fig. 2D; Olson 1999). A recessus conicalis is not present in Anhimidae, and is shallow in *Anseranas*, *Conflicto* (Tambussi et al. 2019), *Nettapterornis* (contra Olson 1999; pers. obs.), and *Presbyornis* (Ericson 1997).

*Asteriornis* exhibits two deep and distinct articular cotyles (Fig. 1 dorsal; Fig. 3D) on both the left and the retrodeformed right articular. This bicotylar condition characterizes Galloanserae (Weber & Hesse 1995; Ericson 1997; Mayr et al. 2018) and *Ichthyornis*, and is markedly distinct from the three-cotyle morphology of palaeognaths and most neoavians. Although the preservation of the lateral cotyle is poor and its total extension cannot be confidently determined, both cotyles are arranged diagonally, with the medial cotyle being rostromedially oriented (Fig. 1). This condition is similar to that of anhimids, *Anseranas*, megapodes, and *Ichthyornis*, and contrasts with the arrangement in Anatidae and non-megapode galliforms, where both cotyles are aligned with the mandibular ramus (Fig. 3D).

**Figure 3.**
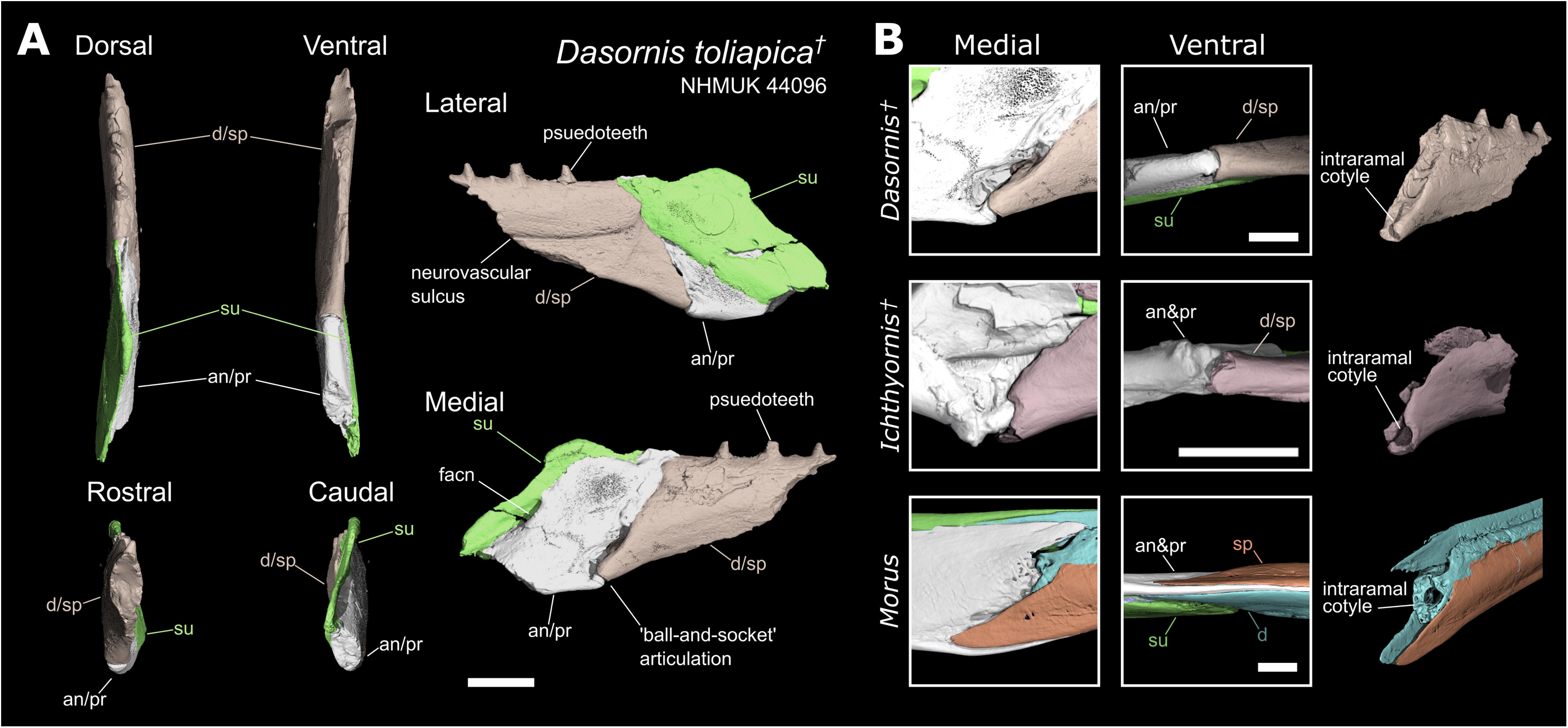
Digitally reconstructed mandible of *Vegavis* and anatomical comparisons with *Ichthyornis* and crown-birds. A) Digitally reconstruction of the left mandibular ramus of *Vegavis* in lateral, medial and dorsal views, from digital meshes of the left caudal mandible and right rostral mandible (mirrored) of specimen AMNH FARB 30899. B) Detailed view of the articular region of the mandible of *Vegavis* and select adult anseriforms in medial view. The broken region not preserving the medial process in *Vegavis* is highlighted in red. Medial processes of the figured anseriforms have been digitally removed and similarly highlighted. C) Isolated post-dentary complexes of the fossil taxa *Vegavis* (AMNH FARB 30899) and *Ichthyornis* (KUVP 119673) as preserved, and digitally isolated in *Anser* and *Morus* in lateral view. D) Dorsal view of the articular region of the fossil taxa *Vegavis*, *Ichthyornis, Asteriornis* and an unidentified Miocene pelagornithid from Peru, and examples of adult galloanserans, a palaeognath and a neoavian. The medial, lateral and caudal cotyles are highlighted and the pelagornithid caudal fossa (which does not articulate with the quadrate) is outlined. Scale bars equal 10mm.

The medial mandibular process of *Asteriornis* is large relative to the overall size of the mandible and is relatively narrow dorsoventrally (Fig. 1 caudal). This is similar to the morphology seen in galloanserans (Fig. 2C-E) and distinctly unlike that of palaeognaths, (Fig. 2B) in which the medial mandibular process projects less prominently from a more bulbous articular region and is dorsoventrally deeper. The precise orientation of the medial mandibular process of *Asteriornis* is unclear because, as preserved in their displaced and distorted positions, the left and right medial mandibular processes are asymmetric (leading to the initial misidentification of the right medial process as a retroarticular process (Field et al. 2020)). The left medial mandibular process has a caudally deflected tip and thus, as observed by Field et al. (2020), is reminiscent of the morphology in extant anseriforms. By contrast, the tip of the right medial mandibular process shows a slight rostrally-oriented, hooked tip. We interpret the right medial mandibular process as likely capturing the original morphology of this structure more closely than the left medial mandibular process, due to the superior preservation of its rostral surface. Moreover, the right medial mandibular process does not show a morphology directly comparable to any of the extant taxa sampled here, instead bearing strongest resemblance to the Palaeocene total-group anseriform *Conflicto antarcticus* (Tambussi et al. 2019; Field et al. 2020).

### Vegavis

The post-dentary complex represents the only overlapping material between specimens MACN-PV 19.748 (referred to *Vegavis* based on its postcranial skeletal morphology) and AMNH FARB 30899 (consisting exclusively of skull material). As reported by Torres et al. (2025), the two mandibles exhibit differences in the shapes of their mandibular cotyles, the presence of deep fossae near the coronoid process in AMNH FARB 30899, and the shape of the ventral margin, with AMNH FARB 30899 showing a convex bulge ventral to the medial cotyle while MACN-PV 19.748 exhibits a straight or shallowly concave ventral margin. However, both specimens share a similar general morphology, including features such as a marked lateral crest, a shallow but distinct fossa in the lateral cotyle, and a retroarticular fossa, features absent in any other known Mesozoic birds. Other shared traits include a medial cotyle approaching the lateral margin, a concave and excavated medial surface, and an extremely thin and dorsoventrally low base of the medial process (Fig. 3A), as well as taphonomic similarities, with both specimens broken at a similar point along the ventral margin. Although these morphological differences raise some doubts as to whether both specimens belong to the same species, the number of shared similarities otherwise unrecognised in other Mesozoic birds point towards both specimens at least representing closely related taxa. While the attribution of AMNH FARB 30899 to *Vegavis iaai* warrants further investigation, we treat it as conspecific with MACN-PV 19.748 for the purposes of this study, in line with Torres et al. (2025).

The currently available CT data for AMNH FARB 30899 (Torres et al. 2025) precludes the differentiation of individual elements in its post-dentary complex, mainly due to the presence of high radiodensity inclusions and poor contrast. As such, we were unable to examine the internal arrangement of its post-dentary elements to the same extent as the other fossil taxa included in this study.

#### Inferences from taphonomy

The mandibular rami of AMNH FARB 30899 are preserved *in situ* and are broken at roughly the same point on both sides, with the left side preserving the post-dentary region and an impression of the dentary/splenial and the right side preserving the reverse. Although the mandibular symphysis is not preserved on either specimen, an impression of the symphysial region is preserved in AMNH FARB 30899. It remains unclear whether the mandibular symphysis was fused or unfused (Torres et al. 2025).

The rostral projection of the probable surangular of the left caudal mandible of AMNH FARB 30899 is well-preserved and appears almost complete, tapering at its rostral end as is typical for the rostral process of the surangular in other birds (Fig. 3C). In a reconstruction combining the left caudal and right rostral portions of the AMNH FARB 30899 mandible, this rostral projection appears to insert into the portion of the mandible comprising the dentary/splenial, as in other avian taxa (see Fig. 2). The interpretation of this process as a projection of the surangular is supported by its relatively narrow mediolateral width relative to the overlapping portion of the right mandibular ramus.

The preservation of the relatively delicate surangular process without associated material from the dentary/splenial components is remarkably similar to known isolated post-dentary complexes of *Ichthyornis* (e.g., KUVP 119673; Fig. 3C), suggesting a lack of fusion between the post-dentary complex (especially the surangular) and the dentary/splenial. Although both known *Vegavis* post- dentary complexes have incomplete rostral ends, it is notable that the ventral portions of both specimens are broken at very similar positions (see Torres et al. (2025) Extended Data Fig. 4). Additionally, the right mandibular ramus of AMNH FARB 30899 is broken at an almost identical position, preserving instead the dentary/splenial rostral to the post-dentary complex. We hypothesise that this is indicative of a structural weakness in that region of the mandible, consistent with the presence of an intraramal joint in *Vegavis*. If the position of this structural weakness represents the rostral extent of the angular bone, it would be suggestive of the presence of a joint similar to the presumably synovial intraramal joint of *Ichthyornis* (Fig. 2A; Fig. 4B) and Pelagornithidae (Fig. 4B). A rostrocaudally short angular would result in little overlap with the dentary/splenial, forming a hinge where the mandible would be likely to separate taphonomically, as is frequently the case in *Ichthyornis* (e.g., KUVP 119673; BHI 6421; FHSU 18702; Supplementary Fig. 2; Field et al. 2018) and in Pelagornithidae (e.g. Mayr & Rubilar-Rogers 2010; Mayr et al. 2021). This contrasts with the condition observed in extant neoavians bearing intraramal joints (e.g., members of Diomedeidae, Fregatidae, Phalacrocoracidae, Sulidae, Pelecanidae, Caprimulgidae and Nyctibiidae (Bühler 1970; Zusi & Warheit 1992)), as well as galloanserans, which exhibit a broad region of insertion of the angular into the dentary/splenial (Fig. 2C-F).

**Figure 4.**
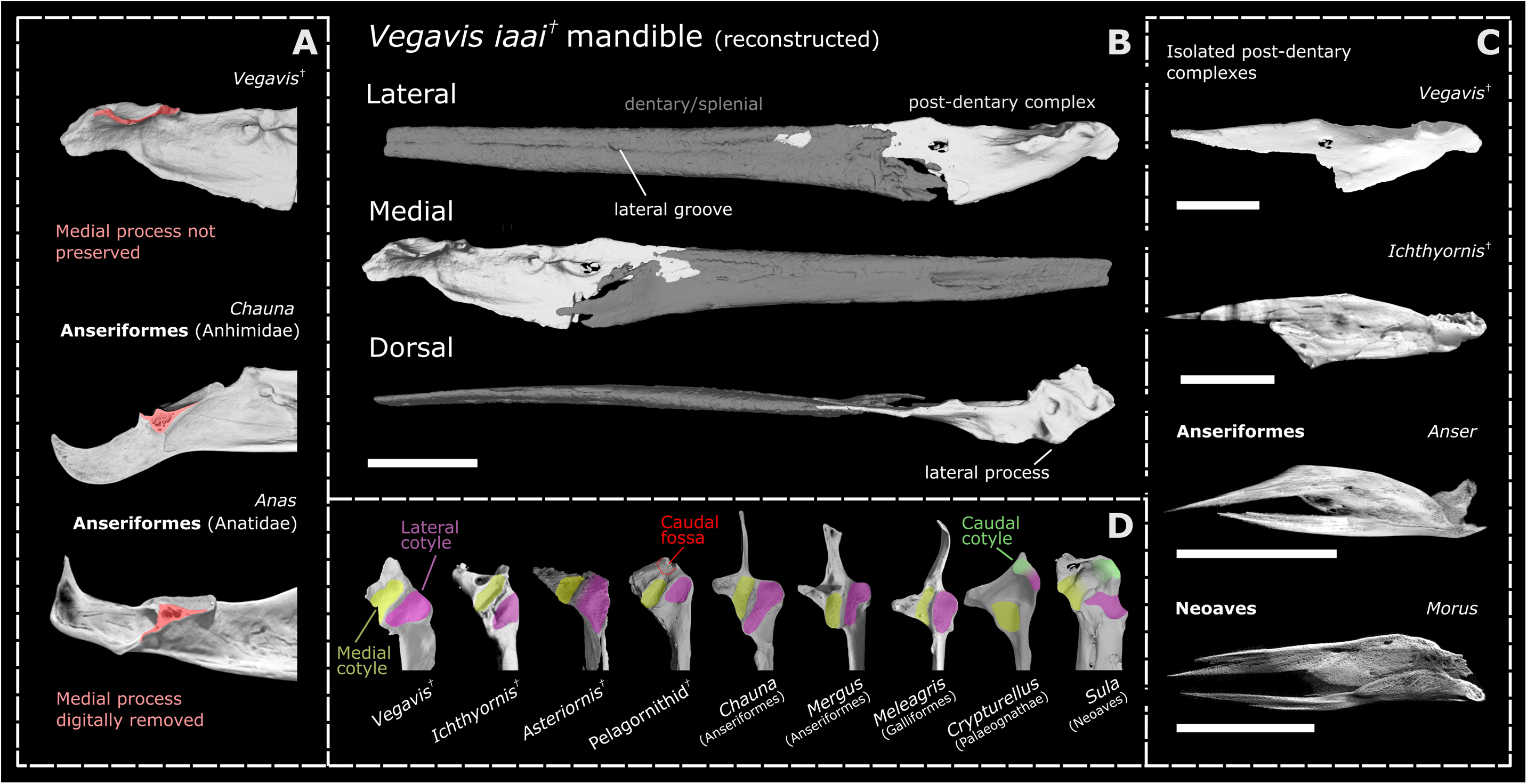
Partial mandible of *Dasornis* (Pelagornithidae) and intraramal joint anatomy. A) Fragment from left ramus in dorsal, ventral, medial, lateral, rostral and caudal views. The angular and prearticular could not be differentiated. As in *Ichthyornis*, the dentary and splenial could not be distinguished and thus are figured with an intermediate colour. Additional abbreviations: facn, fossa aditus canalis neurovascularis. Scale bar equals 10mm. B) Closer view of the intraramal joint anatomy of *Dasornis, Ichthyornis* and *Morus* in medial and ventral view. Isolated dentary/splenial portions of the mandible shown in oblique view, illustrating the rostral articulation of the intraramal joint. Scale bars equal 5mm.

#### General Morphology

The mandibular ramus is straight along its entire length, with the rostral end showing no indication of curving into a rounded mandibular symphysis as in anseriforms and galliforms. The preserved impression of the mandibular symphysis appears to show a narrow, pointed and trough-like morphology, unlike the dish-like shape of even the narrowest anseriform symphyses (e.g., *Mergus*). The rostralmost preserved part of the ramus remains directly dorsoventrally vertical, not showing any sign of the mediolateral slanting which leads into the curved mandibular symphysis in palaeognaths and galloanserans. This narrow vertical portion of the mandibular tip is similar to that of *Ichthyornis* (Fig. 2A), pelagornithids, and some neoavians with intraramal hinges (e.g., *Morus*; Fig. 2F).

The dorsal edge of the ramus is straight along most of its length, lacking the dorsoventral curvature seen in palaeognaths, galliforms and anhimids. Whilst the straight profile of the dentary/splenial region resembles that of some anatids (e.g., Fig. 2D), the post-dentary complex does not exhibit the prominent mandibular apex seen in this group.

The ventral margin of the dentary/splenial is curved such that the ramus increases in dorsoventral depth, reaching a maximum ventral expansion at the dentary/post-dentary contact. The overall profile of the mandible is distinctly unlike that of known anseriforms, which exhibit dorsoventrally straight or slightly concave ventral margins and variably convex dorsal margins. Even anseriforms with narrow and pointed mandibles most comparable to *Vegavis* (e.g., *Mergus*) retain a prominent mandibular apex and minimal ventral expansion, resulting in a strikingly different overall profile. The condition in *Vegavis* more closely resembles the overall mandibular shape of *Ichthyornis* (Fig. 2A), pelagornithids, and some piscivorous neoavians (e.g., *Morus* Fig. 2F; *Gavia* Torres et al. 2025), yet it differs from these extinct taxa in the position of the articular region, which aligns with the dorsal margin of the mandible rather than being closer to the ventral margin.

#### Splenial/Dentary

The caudal end of the dentary/splenial region is not well-preserved, so the degree to which the dentary is caudally forked into dorsal and ventral processes is uncertain; as preserved, it appears either unforked (as in Anatidae, *Ichthyornis* and Pelagornithidae) or asymmetrically forked with a small dorsal fork (as in galliforms, Anhimidae and *Asteriornis*). The lateral surface exhibits a prominent groove running rostrocaudally along the midline resembling the lateral grooves of pelagornithids (neurovascular sulcus; Mayr & Rubilar-Rogers 2010), *Ichthyornis* (Fig. 2A) and the piscivorous neoavians *Gavia* and *Phaethon*. *Vegavis* does not exhibit a lateral mandibular ridge like that of Anatidae (Fig. 2D).

#### Post-Dentary Complex

Ventral to the rostrally pointed surangular described above, the rostral margin of the post-dentary complex is dorsoventrally vertical and almost straight, with a mostly flat anterior surface and a slightly rounded ventral end probably formed by the angular, forming a probable articular surface consistent with the presence of an intraramal joint (Fig. 2C). As discussed above, the nature of this probable joint is highly reminiscent of those in *Ichthyornis* and the pelagornithid *Dasornis*, though in these taxa the anterior margin of the post-dentary complex is dorsoventrally sloped (Fig. 4B), rather than vertical as in *Vegavis*. Furthermore, while both *Ichthyornis* and *Dasornis* exhibit clear articular condyles on the caudoventral margin of the intrarramal joint (Fig. 4B), this region is flat or slightly rounded in *Vegavis*, though condyles are also absent in derived pelagornithids such as *Pelagornis chilensis* and MUSM 1677 (see discussion below). The morphology of this region in *Vegavis* contrasts markedly with that of neoavians with intraramal hinges, in which the angular extends into a long sheet-like rostral projection inserting between the splenial and dentary, rather than forming a rounded condyle (Fig. 4B, Supplementary Fig. 4).

Although no complete medial mandibular process of *Vegavis* is preserved, inferences pertaining to its general morphology can be made based on preserved portions of the articular region. The medial edge of the preserved articular region, including the margin of the medial cotyle, is extremely narrow dorsoventrally (Fig 3B). The preserved broken edge exhibits only a fine ridge of bone delimiting the ventral portion of the medial cotyle. This morphology contrasts with that of extant anseriforms, in which the ridge of bone lining the rostral edge of the medial cotyle is similarly thin, but becomes more robust and subtriangular in cross-section at the caudal part of the cotyle. This more robust cross-section is partly due to the presence of a distinct ridge on its ventral apex, defining a clear internal cavity. This forms the base of the characteristic elongate, dorsally orientated medial mandibular process of anseriforms (Fig. 3A). *Vegavis* does not exhibit a subtriangular expansion at the base of the medial mandibular process as in anseriforms, and we can confidently infer that it would not have exhibited an anseriform-like medial mandibular process. The medial surface of the articular region appears to have instead been demarcated by a thin ridge of bone delimiting the medial cotyle and a small medial process, akin to the morphology seen in some piscivorous neoavians such as pelecaniforms, and unlike the more robust medial processes of *Ichthyornis* and pelagornithids (Fig. 3D).

The well-preserved caudal ends of the mandibular rami in both AMNH FARB 30899 and MACN-PV 19.748 do not exhibit a recessus conicalis (found in Anseres and several fossil total-clade Anseriformes; Fig. 2D). The lateral mandibular process is small and indistinct relative to the large ‘lateral crest’ (Álvarez-Herrera et al. 2024) rostral to it. The two structures are separated by a small depression, resembling that of some neoavians such as Phalacrocoracidae and Gaviidae. A fossa ventral to the medial tubercule likely represents the fossa aditus canalis neurovascularis, a structure also seen in pelagornithids, anseriforms, and neoavian birds such as *Gavia* (Baumel & Witmer 1993).

The mandible of *Vegavis* exhibits two mandibular cotyles (Fig. 3D), forming a bicondylar articulation with the quadrate as in Galloanserae, some neoavian taxa (e.g., Columbidae), *Ichthyornis*, and Pelagornithidae (which exhibit an additional caudal fossa, which has been referred to as a caudal cotyle (Mayr & Rubilar-Rogers 2010) although it does not articulate with the quadrate). As noted by Torres et al. (2025), the two mandibular cotyles are positioned differently relative to those of extant galloanserans. The galloanseran condition has two cotyles which are approximately mediolaterally adjacent. In anatids and galliforms these cotyles are parallel (i.e. craniocaudally aligned with the main axis of the mandibular ramus), whereas in Anhimidae, Megapodiidae and *Asteriornis* these are diagonally arranged, with the medial cotyle more caudally positioned (Fig. 3D). By contrast, *Vegavis* exhibits a medial cotyle positioned caudomedial to the lateral cotyle, angled obliquely relative to the main axis of the mandibular ramus such that the lateral portion of the medial cotyle is directly caudal to the medial part of the lateral cotyle, and both cotyles are almost perpendicular to the mandibular ramus (Fig. 3D). The position of these cotyles in *Ichthyornis* and pelagornithids appears intermediate between the condition seen in Galloanserae and that of *Vegavis* (Fig. 3D), with the medial and lateral cotyles rostrocaudally offset such that the lateral portion of the medial cotyle is directly caudal to the medial part of the lateral cotyle. The cotyles of *Ichthyornis*, however, exhibit a greater degree of mediolateral overlap than in *Vegavis*.

### *Dasornis* (Pelagornithidae)

The left mandible fragment of the *Dasornis toliapca* holotype comprises only a small portion of the mandibular ramus surrounding the intraramal joint, the general anatomy of which was initially described by Owen (1873). Due to pyrite infill in this fossil noted by Milner and Walsh (2009), the internal anatomy of this mandible is difficult to assess using CT-scans, and some anatomical details enumerated below should be treated as hypotheses to be tested with superior material should any come to light. Our inferences are supported with reference to published pelagornithid mandible material (e.g., *Pelagornis chilensis* (Mayr & Rubilar-Rogers 2010)*, Lutetodontopteryx tethyensis* (Mayr & Zvonok 2012), *Protodontopteryx ruthae* (Mayr et al. 2021)) and surface scans of MUSM 1677.

#### Dentary/Splenial

The dentary and splenial could not be distinguished from one-another (Fig. 4A). This could be due to poor preservation obscuring any sutures but, given the detailed preservation of much of the external mandibular surface, we consider it probable that the suture between these elements was developmentally obliterated through fusion, with both elements being similarly undifferentiable in other pelagornithids (e.g. Mayr & Rubilar-Rogers 2010). This is also the condition in *Ichthyornis*, in which the dentary and splenial can only be differentiated along a short caudally positioned section in some specimens (e.g. YPM 1450, BHI 6421, Supplementary Fig. 8) and are otherwise completely fused, even in skeletally immature individuals. This fused condition contrasts with that of adult neoavians with intraramal joints (including members of Diomedeidae, Fregatidae, Phalacrocoracidae, Sulidae, Phaethontidae and Pelecanidae), which show a visibly distinct suture between the dentary and splenial retained into skeletal maturity (Zusi & Warheit 1992) (Supplementary Fig. 4).

The dentary/splenial portion of the mandibular ramus is robust and slightly mediolaterally wider than the post-dentary complex (Fig. 4A dorsal). This is similar to the condition in *Ichthyornis* (Fig. 2A), in which the dentary/splenial region is wider and more robust than the post-dentary complex. A prominent neurovascular sulcus (Mayr & Rubilar-Rogers 2010) runs along the lateral surface of the dentary-splenial complex, which exhibits a slight dorsal deflection at its caudal end (Fig. 4 lateral). A similar groove is also present on the lateral surface in *Vegavis* and *Ichthyornis*, although in both taxa the groove is broader and less defined towards the caudal end of the dentary/splenial than in *Dasornis*. Amongst extant taxa, similar grooves are present in *Phaethon* and *Gavia*.

The caudal end of the dentary/splenial complex is unforked and slopes steeply towards the ventral edge of the mandible. An unforked dentary is a condition shared with Anseres (Fig. 2D), *Ichthyornis* (Fig. 2A) and potentially *Vegavis* (Fig. 3B).

#### Post-dentary complex

The rostral termination of the surangular is mediolaterally narrow and restricted to the dorsal portion of the ramus, inserting into the dentary/splenial complex (Fig. 4 dorsal, lateral). The surangular continues rostrally past the intraramal joint and persists beyond the broken caudal extent of this specimen. The ventral margin of the surangular slopes ventrally caudal to the intraramal joint, and the surangular is mediolaterally narrow relative to its dorsoventral depth.

Aside from the approximate margin of the surangular, sutures among the remaining osteological components of the post-dentary complex could not be clearly differentiated (Fig. 4). The rest of the preserved post-dentary complex most likely consists of portions of both the angular and prearticular. By comparison to the better-known mandibular morphology of other birds (Fig. 2), the angular most likely comprises the ventral portion of the post-dentary complex. This includes a bulbous process or condyle at the rostroventral margin of the angular/prearticular which inserts into a socket formed by the dentary/splenial (Fig. 4B; see Intraramal Joint below). The dorsal extent of the undifferentiated post-dentary material suggests that the prearticular extends rostrally to meet the intraramal joint (Fig. 4 medial). This can be surmised because the undifferentiated elements caudal to the intraramal joint extend to the joint’s dorsal margin, which is a position occupied by the prearticular in all observed taxa in which this element could be distinguished (Fig. 2). The rostral projection of the prearticular is comparable to that of *Ichthyornis* (Fig. 2A), anseriforms (Fig. 2C,D), and neoavians with intraramal joints (Fig. 2F). Given the rostral extent and assumed dorsal position of the prearticular it can be inferred that this element comprises the ventral margin of the fossa aditus canalis neurovascularis (Baumel & Witmer 1993) as in some anseriforms such as *Anser* (Supplementary Fig. 9). Additionally, the inferred dorsal margin of the prearticular exhibits a sharp ventrally sloping edge caudal to the intraramal joint. This is most closely reminiscent of the *Ichthyornis* prearticular (Fig. 2A), althought the prearticulars of anseriforms and neoavians with intraramal hinges can also exhibit ventrally sloping edges, albeit at a less steep angle (Fig. 2D,F).

#### Intraramal joint

The splenial/dentary is clearly differentiated from the post-dentary complex at the intraramal joint. The caudal end of the dentary/splenial is characterised by a deep and concave cotyle on its ventral margin, bounded by raised lateral and medial edges (Fig. 4B). As described above, a matching round condyle is developed on the rostroventral margin of the angular/prearticular, forming a likely synovial joint. This synovial joint is present in other pelagornithids (Zusi & Warheit 1992; Mayr & Rubilar-Rogers 2010, Mayr & Zvonok 2012, Mayr et al. 2021); however, a well-defined condyle in the angular/prearticular appears to be absent in *Pelagornis chilensis* (Mayr & Rubilar-Rogers 2010) and MUSM 1677, with a subtle, less distinct condyle present in *Pelagornis sandersi*. This ‘ball and socket’ type of articulation is strikingly similar to the condition in *Ichthyornis*, which also exhibits a deep cotyle in the dentary/splenial and a round condyle on the angular (Fig. 4B). However, the cotyle of *Ichthyornis* is deeper with a taller lateral margin, and, unlike in *Dasornis,* the condyle is situated slightly rostral to the ventral tip of the angular. This condylar morphology of the angular contrasts with the condition of extant neoavians (Fig. 2F), in which the angular is mediolaterally narrow rather than bulbous, and fits into a deep insertion between the dentary and splenial (Fig 4B, Zusi & Warheit 1992). A condylar articulation is also developed convergently in some members of Neoaves, most clearly exemplified by *Morus* (Fig. 4B, Supplementary Fig. 4), but also present in some other piscivorous taxa including Fregatidae, Phalacrocoracidae, and Phaethontidae. However, in Neoaves this articulation is formed by a novel post-hatching ossification of the Meckel’s cartilage spanning the intraramal hinge (the “internal ossification”; Zusi & Warheit 1992), which can form a highly variable and complex multi-jointed articulation. In *Morus* its rostral portion forms a deep rounded cotyle, incorporated into the dentary, and its caudal portion forms a complex three-pronged condyle, incorporated into the prearticular. As such, the neoavian synovial joint, when present, appears to represent a novel structure situated entirely dorsal to the sheet-like rostral projection of the angular, instead of forming the ventral margin of the intraramal hinge as in *Dasornis, Ichthyornis* and likely *Vegavis*.

Further dorsally a small gap exists between the dentary/splenial and the post-dentary complex both in *Dasornis* and *Ichthyornis*. Along the dorsal half of the joint, the surangular inserts into the splenial/dentary via a narrow rostral process (Fig. 4, dorsal). The rest of the post-dentary complex, most likely incorporating the prearticular, inserts weakly into a shallow groove or depression on the caudal end of the dentary/splenial; a similar articulation is present in *Ichthyornis*, yet this groove is deeper and bounded by a higher lateral margin (Fig. 2A, 4B). In contrast, in neoavians with intraramal hinges the prearticular overlies the medial surface of the dentary, and no similar articular groove is developed. In these taxa the internal ossification is incorporated into the dentary and forms a discrete, rounded cotyle for the condyle of the prearticular (Fig. 2F, 4B).

## Discussion

### Asteriornis

#### An early neornithine with some unique mandibular features

The mandible of *Asteriornis* is clearly distinct from that of crownward stem-birds such as *Ichthyornis* in its edentulism, fusion of the mandibular symphysis, and lack of a predentary bone. In addition to these features shared with all extant birds, *Asteriornis* exhibits some apparently autapomorphic characteristics in its lower jaw: its mandibular rami are narrower mediolaterally and deeper dorsoventrally than other taxa examined, its mandibular symphysis exhibits a distinctly ‘scooped’ morphology, and its surangular does not appear to contact the angular caudally. The dorsoventrally and rostrocaudally deep, convex mandibular symphysis bears resemblance to a controversial small isolated mandibular symphysis from the Upper Cretaceous Lance Formation with a comparable morphology, initially identified as an improbably early representative of Psittaciformes (Stidham 1998; Hope 2002), but later regarded as potentially belonging to the edentulous non-avialan theropod clade Caenagnathidae (Dyke & Mayr 1999; Waterhouse 2006; Mayr 2022). However, the Lance Formation specimen lacks the distinctive mandibular grooves of Caenagnathidae, which appear early in development and are present even in the smallest members of the clade (Funston et al. 2020). As such, it is plausible that the Lance Formation specimen could instead belong to an *Asteriornis*-like bird (albeit one considerably larger than *Asteriornis* itself). The presence of *Asteriornis*-like birds in the latest Cretaceous of North America might be suggested by the recent description of the partial skeleton of an immature bird from the Lance Formation preserving a quadrate strikingly similar to that of *Asteriornis* (Brownstein 2024). Further investigation and future fossil discoveries may help to assess the extent to which the seemingly autapomorphic features of the *Asteriornis* mandible are distinct from this taxon.

#### Does Asteriornis have a palaeognath-like mandible?

Torres et al. (2021) recovered *Asteriornis* among total-clade palaeognaths, although that phylogenetic analysis was focused primarily on Cretaceous stem birds. Here, we demonstrate a lack of derived features shared between *Asteriornis* and palaeognaths; many of the traits we identify as being shared between the mandibles of *Asteriornis* and palaeognaths (e.g., a lack of contact between the splenial and prearticular, a relatively small splenial, and a straight dorsal margin of the surangular without a prominent mandibular apex) are also shared with galliforms. Rather than representing apomorphic traits uniting *Asteriornis* with palaeognaths, these morphological features may therefore represent crown bird symplesiomorphies or convergent morphological similarities between galliforms and palaeognaths.

The type specimen of *Asteriornis* appears to have been either skeletally mature or approaching skeletal maturity, on the basis of osteohistology of the femur (Field et al. 2024), and as such the limited fusion of the rostral elements of its skull is not necessarily indicative of osteological immaturity. However, caution must be taken as cranial elements are often among the latest to fuse over avian ontogeny, and cranial sutures may remain open in otherwise skeletally mature birds (Arnaout et al. 2025)). Weakly fused rostral skull elements are typical of adult galliform birds (Field et al. 2020), and similarly, visible sutures between the dentary, angular and surangular at osteological maturity are known in galliforms (Hogg 1977; 1983), so the ability to distinguish among these bones in an osteologically mature bird is not unexpected. The complete lack of fusion between the splenial and the rest of the mandibular ramus (to the point of disarticulation in the right ramus) is unusual, and represents one of the more compelling morphological features shared between *Asteriornis* and palaeognaths. This characteristic is, however, also shared with the galliform subclade Megapodiidae (Field et al. 2020), the sister taxon to the remainder of Galliformes, and thus does not unambiguously unite *Asteriornis* with palaeognaths to the exclusion of galloanserans.

Torres et al. (2021) reported a single synapomorphy supporting the position of *Asteriornis* as a stem-palaeognath: a deeply forked caudal end of the dentary with dorsal and ventral forks of approximately equal length. While the dataset yielding this result only included a limited sample of crown birds and was thus suboptimal for assessing the affinities of *Asteriornis* (Benito et al. 2022b), the present work also reveals that the caudal end of the *Asteriornis* dentary in fact exhibits a short and indistinct dorsal fork more similar to that of extant galliforms and anhimids (Fig. 1 right lateral). Given this re-interpretation of the only putative synapomorphy of an exclusive *Asteriornis* + Palaeognathae clade, we consider the position of *Asteriornis* as a stem-palaeognath highly unlikely.

#### Asteriornis—the case for galloanseran affinities based on mandibular morphology

Despite the absence of direct evidence of large, hooked retroarticular processes as originally hypothesised (Field et al. 2020), the mandible of *Asteriornis* shows many previously unrecognised morphological similarities with those of extant galloanserans. Although potentially plesiomorphic for Neornithes, the presence of two mandibular cotyles is congruent with galloanseran affinities (Weber & Hesse 1995; Ericson 1997; Mayr et al. 2018). Other traits shared with galloanserans include a more caudally positioned splenial relative to that of palaeognaths, and the presence of a large, dorsoventrally narrow medial mandibular process of the articular.

The taphonomic loss of the caudal extremity of the holotype mandibles renders it impossible to confirm the presence of galloanseran-like retroarticular processes in *Asteriornis,* yet the morphology of the preserved caudalmost region of the post-dentary complex is remarkably similar to that of galloanserans, especially galliforms, and quite distinct from that of taxa lacking retroarticular processes such as palaeognaths and most neoavians (Crane et al. 2025). The potential presence of this key galloanseran synapomorphy would distinguish *Asteriornis* from other putative total-clade galloanserans for which its absence is known, including *Vegavis* and Pelagornithidae, thus additional mandibular material from *Asteriornis* will be invaluable (Mayr & Rubilar-Rogers 2010; Clarke et al. 2016; Mayr et al. 2018; Torres et al. 2025).

#### Wonderchicken or Wonderduck?

Field et al. (2020) suggested that *Asteriornis* exhibited a unique combination of galliform-like and anseriform-like features. We find that this holds true for the mandible to only a limited extent; whilst certain aspects of the mandible resemble those of anseriforms, we find the lower jaw of *Asteriornis* to be overwhelmingly galliform-like. Galliform-like traits of the mandible shared with *Asteriornis* include the narrow cross-sectional profile of the mandibular rami, a relatively small, rod-shaped splenial and a surangular with a straight dorsal margin and forms a shallow lateral process. The caudal end of the dentary from our 3D reconstruction is especially galliform-like due to the highly asymmetric lengths of the dorsal and ventral processes, as opposed to the deeply forked nature of the palaeognath mandible or the unforked mandible of Anseres. Some of these galliform-like features of *Asteriornis* are also shared with the early-diverging anseriform group Anhimidae, most notably a straight dorsal margin without a prominent mandibular apex and the asymetrically forked dentary.

In its initial description, the presence of dorsoventrally deep and hooked ‘retroarticular processes’ was considered one of the key anseriform-like features of *Asteriornis*; the reinterpretation of this retroarticular process as a displaced medial mandibular process (Crane et al. 2025) necessitates a reassessment of the extent to which the mandible of *Asteriornis* resembles that of anseriforms. Other anseriform-like features of the mandible of *Asteriornis* include rostrocaudally straight mandibular rami without concave ventral curvature (as in Anatidae) and a rostrocaudally extensive prearticular. The straight mandible without ventral curvature is also exhibited by the possible stem-anseriform *Anachronornis* (Houde et al. 2023) and other fossil total-group anseriforms such as *Conflicto* and *Paakniwatavis* (Tambussi et al. 2019; Musser & Clarke 2024), which also show anatid-like prominent mandibular apexes. The question as to whether the galliform-like mandible of anhimids is plesiomorphic for anseriforms or derived from an ancestrally more anatid-like mandible remains uncertain (Houde et al. 2023). The prevalence of a straight mandibular morphology combined with a prominent mandibular apex in fossil total-group anseriforms suggests that this condition is likely ancestral for the group, but other aspects of the anatid mandible (e.g., spatulate mandibular symphysis, recessus conicalis) are likely derived. It is possible that the ancestral anseriform mandible exhibited a mosaic of anatid-like and anhimid-like traits.

Several anseriform-like traits of the *Asteriornis* mandible are also shared with *Ichthyornis* and might constitute crown bird symplesiomorphies rather than anseriform synapomorphies; as such, our observations provide little discrete character support for an exclusive *Asteriornis* + Anseriformes clade (*contra* the proposal by Mayr 2022). Additionally, we interpret the medial mandibular process as likely having exhibited a rostrally deflected tip based on the better-preserved morphology of the right medial mandibular process. This is unlike the anseriform-like, caudally deflected morphology initially interpreted by Field et al. (2020) based on the incomplete left medial mandibular process. The rostrally deflected tip of the medial mandibular process is, however, like that of the fossil total-group anseriforms *Conflicto* and *Anachronornis*, representing a previously unrecognised feature associated with total-clade Anseriformes. Overall, however, there are only a limited number of morphological similarities between the mandible of *Asteriornis* and that of anseriforms.

The more galliform-like mandible of *Asteriornis* could be indicative of a phylogenetic position closer to crown Galliformes than to crown Anseriformes. This would support the position recovered using Bayesian methods by Field et al. (2020), and both Bayesian and parsimony methods in Crane et al. (2025), which recovered *Asteriornis* as a stem-galliform. An alternative interpretation could be that features of the mandible here considered as typically galliform could instead represent galloanseran symplesiomorphies, with the galliform mandible more closely resembling the ancestral galloanseran condition than that of anseriforms. This is supported by the shared morphological traits of *Asteriornis*, galliforms, and anhimid anseriforms, suggesting that the ancestral galloanseran may have also exhibited features such as a forked dentary (although, as discussed previously, some aspects of the galliform-like anhimid mandible may have been independently derived). This galliform-like ancestral condition would be in line with the galliform-like quadrates of the total-group anseriforms *Presbyornis* and *Conflicto*, which are also strikingly similar to that of *Asteriornis* (Kuo et al. 2023), as well as recent interpretations of galloanseran cranial form from investigations integrating developmental and palaeontological data (Arnaout et al. in prep).

### Vegavis

#### Does Vegavis have an anseriform-like mandible?

*Vegavis* has been recovered as a total-clade representative of Anseriformes in several phylogenetic analyses, both as a crown-group anseriform (Clarke et al. 2005; Clarke et al. 2016; Musser and Clarke 2024; Torres et al. 2025) and as a stem-group anseriform (Worthy et al. 2017). We find the mandible of *Vegavis* to lack any features unambiguously linking it to Anseriformes, such as the rostrocaudally running ridge on the lateral surface of the anatid dentary, or the recessus conicalis, which is present in extant Anseres and most fossil total-clade anseriforms such as *Presbyornis, Conflicto,* and *Nettapterornis* (Ericson 1997; Tambussi et al. 2019; Houde et al. 2023). With currently available material and CT data it is not possible to assess whether *Vegavis* exhibits evidence of other mandibular synapomorphies of Anseriformes, such as a large, triangular splenial or a rostrally- extensive prearticular. The general morphology of the *Vegavis* mandible (Torres et al. 2025) is uncurved dorsoventrally (as in Anatidae) but lacks the prominent mandibular apex of anatids. Otherwise, the overall shape of the mandible is very unlike that of most extant anseriforms, with the narrow, dorsoventrally-oriented rostral end of the preserved dentary/splenial showing no indication of forming the spatulate mandibular symphysis emblematic of most of this clade.

Furthermore, evidence of galloanseran affinities in the mandibular morphology of *Vegavis* is similarly scarce. We interpret the mandible as exhibiting two articular cotyles (contra Álvarez-Herrera et al. 2024) but, as discussed by Torres et al. (2025), note that the positions of the two cotyles are unlike those of extant Galloanserae. Aside from the bicondylar articulation of the articular and quadrate (which is also present in *Ichthyornis* and likely represents a crown bird symplesiomorphy), *Vegavis* lacks most diagnostic galloanseran mandibular traits. Most notably, in contrast to earlier interpretations based on less complete mandibular remains (e.g., Agnolín et al. 2017), *Vegavis* does not exhibit the enlarged and hooked retroarticular process characteristic of all extant Galloanserae. Additionally, the preserved articular region suggests that it is unlikely that *Vegavis* exhibited a large, robust and prominent galloanseran-like medial mandibular process (Fig. 2C-E; Fig. 3A). Furthermore, the overall shape of the mandibular ramus in *Vegavis* is unlike that of any extant galloanseran, with an almost straight dorsal edge and a ventral expansion of the depth of the ramus, peaking at the caudal terminus of the splenial/dentary.

#### Is the Vegavis mandible clearly neornithine?

Phylogenetic analyses do not unanimously unite *Vegavis* with Galloanserae; several analyses have recovered alternative positions for *Vegavis* within Neornithes, or even outside the avian crown group (O’Connor et al. 2011; McLachlan et al. 2017; Field et al. 2020). *Vegavis* is edentulous, as is characteristic of Neornithes, but whether it exhibited a fused mandibular symphysis and lacked a predentary bone, both of which may represent crown bird synapomorphies, remains unknown. Thus, based on currently known specimens, data from the mandible of *Vegavis* provide little evidence regarding its neornithine status.

Importantly, we note previously unrecognised similarities between the mandibles of *Vegavis* and the crownward stem-neornithine *Ichthyornis*, as well as the enigmatic Pelagornithidae. Most notable is the morphology of the post-dentary complex: the preserved region in of the left mandible of AMNH FARB 30899 bears a striking resemblance to the isolated post-dentary complex of *Ichthyornis* (e.g., KUVP 1196730). Other morphological similarities to *Ichthyornis* include a medial articular cotyle positioned caudal to the lateral cotyle (in contrast to the mediolaterally adjacent position of these cotyles in extant Galloanserae) and a rostrocaudally running groove along the midline of the lateral surface of the mandible, also found in in pelagornithids. Furthermore, the overall shape of the *Vegavis* mandible bears a notable similarity to that of both *Ichthyornis* and pelagornithids, including a maximum dorsoventral depth immediately caudal to the dentary/splenial contact and a notable lack of mediolateral curvature along the mandible’s entire preserved length (Mayr & Rubilar-Rogers 2010; Mayr et al. 2021). Some of these features are also present in piscivorous neoavians, including the lateral groove present in Gaviiformes and Phaethontidae, as well as the ventrally expanded mandible of some Suliformes (e.g., *Morus*), and may have evolved convergently.

The nature of the articulation between the dentary/splenial and the post-dentary complex of *Vegavis* is unlike that of any known neoavians, and is strongly reminiscent of that of *Ichthyornis* and pelagornithids. Taphonomic similarities across multiple specimens (MACN-PV 19.748, AMNH FARB 30899) lead us to interpret the mandible of *Vegavis* as exhibiting a low level of fusion between the splenial/dentary and post-dentary complex, possibly indicative of the presence of an intraramal hinge. Multiple groups of extant piscivorous neoavians and extinct toothed (and pseudotoothed) birds exhibit intraramal kinesis as a means of facilitating a wider gape (Martin & Naples 2008). However, in these taxa the angular exhibits a long contact with the dentary laterally and the splenial medially, unlike the simpler dorsoventral articulation apparent from both known *Vegavis* lower jaws. If, as we suggest, the structural weakness at the ventral edge of the post-dentary complex corresponds to the rostral terminus of the angular bone, then this morphology is similar to that of *Ichthyornis* and pelagornithids, which exhibit a simple condylar synovial intraramal articulation, contrasting with the more complex articulation of Neoaves, which can include a synovial joint (e.g., *Morus*), but is primarily formed by long, longitudinally overlapping bone insertions (Zusi & Warheit, 1992).

Overall, congruent with the findings of Mayr et al. (2018) and Álvarez-Herrera et al. (2024), we find no morphologies of the mandible unambiguously uniting *Vegavis* with Anseriformes, Galloanserae, or indeed any other major crown bird lineage. Given the extent to which aspects of galloanseran cranial morphology may be plesiomorphic for crown group birds (with the notable exception of certain features such as the retroarticular process), we consider mandibular morphology to call the position of *Vegavis* as a total-group galloanseran into question. Indeed, morphological similarities with the mandible of the stem-bird *Ichthyornis* lend support to interpretations of a position for *Vegavis* either extremely deep within, or even outside the avian crown-group (O’Connor et al. 2011; McLachlan et al. 2017; Field et al. 2020; Field 2025).

### *Dasornis* (Pelagornithidae)

#### Do pelagornithids have a galloanseran mandible?

Despite most phylogenetic analyses to date having resolved pelagornithids as members of total-group Galloanserae (Bourdon 2005; Mayr 2011; Field et al. 2020; Musser and Clarke 2024), morphological support for this relationship is relatively sparse. A total-clade galloanseran position is primarily supported by a select few morphological traits relating to palatal and mandibular morphology (Mayr 2022) but this evidence has been weakened by the recent suggestion that aspects of galloanseran palatal morphology may be plesiomorphic for Neornithes (Benito et al. 2022a; Benito et al. 2025; Field et al. 2025). Similarly, recent suggestions that a bicondylar articulation of the quadrate with the articular is plesiomorphic for Neornithes rather than representing a galloanseran synapomorphy casts further doubt on the galloanseran affinities of pelagornithids (Mayr & Rubilar-Rogers 2010; Mayr et al. 2018; Kuo et al. 2023). Moreover, one of the strongest synapomorphies of Galloanserae is their large and hooked retroarticular processes, which are present in all living galloanserans and notably absent in pelagornithids (Mayr et al. 2018; Mayr & Rubilar-Rogers 2010). Mandibular support for pelagornithids as members of total-clade Galloanserae therefore appears weak.

Nonetheless, the holotype of *D. toliapica* exhibits some aspects of mandibular morphology that are shared with galloanserans, specifically anseriforms, and are exhibited by other pelagornithid specimens (Mayr & Rubilar-Rogers 2010, Mayr & Zvonok 2012, Mayr et al. 2021). These include a caudally unforked dentary, which is characteristic of anseriform mandibles, and a prominent fossa aditus canalis neurovascularis, which is found in some anseriforms such as *Anser* (Baumel & Witmer 1993). Additionally, we interpret the prearticular to extend rostrally up to the intraramal joint, making it of comparable size to that of anseriforms. However, aside from the fossa aditus canalis neurovascularis, these features are also present in the stem bird *Ichthyornis*, suggesting that these traits are most likely plesiomorphic for Neornithes rather than constituting galloanseran synapomorphies.

#### Do pelagornithids have a neornithine mandible?

The cranial morphology of pelagornithids is unusual amongst crown birds, including an unfused fronto-parietal suture not seen in any other neognath taxa (Mayr 2011). The mandible of pelagornithids is similarly unusual; well-established peculiarities include an unfused mandibular symphysis and the presence of bony pseudoteeth.

If pelagornithids are crown birds they would be unique in exhibiting an unfused mandibular symphysis. This, combined with the presence of an intraramal joint, would have enabled a highly kinetic mandible unlike that of any living bird (Zusi & Warheit 1992). In addition to their unfused symphysis, an underexamined ossification at the rostral tip of the jaws unobserved in any other crown birds has been interpreted as a predentary ossification (Mayr & Rubilar-Rogers 2010; S. Wang et al. 2020), raising the possibility that it is homologous with the predentaries known seen in some non- neornithine Euornithes (and absent in any known crown birds). Nonetheless, the detailed nature and homologies of this element remain in need of deeper investigation in light of a current understanding of the predentary ossification (Bailleul et al. 2019).

Pelagornithids are unique in their possession of bony pseudoteeth—tooth-like projections issuing from the mandible and premaxilla (Louchart and Viriot 2011). These differ from the true teeth of stem-birds in their composition and arrangement: they are made of bone with a keratinised external covering, as opposed to enamel and dentine, and grow directly from the jaw bone rather than inserting into tooth sockets or a communal groove in the dentary (Louchart et al. 2013; Louchart et al. 2018; Brocklehurst & Field, 2021). Despite these differences, recent analyses have proposed that these pseudoteeth are developmentally homologous with true teeth (Mayr & Rubilar-Rogers 2010; Mayr & Zvonok 2012; Louchart et al. 2018), which may be indicative of an early split between pelagornithids and other neornithines (Mayr 2022), and may support a step-wise process of tooth reduction along the crownward portion of the avian stem lineage (Dumont et al. 2016; Wang et al. 2017). Given that all known toothed euornithines also exhibit unfused mandibular symphyses (and thus, all euornithines with fused symphyses have toothless mandibles), the possibility that these traits are developmentally correlated cannot be dismissed. Although not all euornithines with edentulous mandibles have fused mandibular symphyses (e.g., *Archaeorhynchus*, *Mengciusornis*; Wang & Zhou 2017; M. Wang et al. 2020), it is possible that tooth loss is a necessary precursor for evolving a fused symphysis in Euornithes. As such, in a scenario in which pelagornithids might have ‘re-evolved’ tooth-like structures from edentulous ancestors by re-appropriating conserved genetic pathways, these developmental mechanisms might be similarly linked to the loss of a fused symphysis.

Furthermore, we identify additional aspects of pelagornithid mandibular morphology shared with *Ichthyornis* and other stem-birds to the exclusion of neornithines, including a mediolaterally thick and inflexible dentary, and a splenial that is more robust than the post-dentary complex. The dentary and splenial of *D. toliapica* are indistinguishable due to fusion, in contrast to the condition in neoavian birds with intraramal joints, which exhibit a distinct splenial-dentary suture even as adults (Zusi & Warheit 1992). The condition in *D. toliapica*, however, is similar to the highly fused dentary and splenial of *Ichthyornis* (Fig. 2A; Supplementary Fig. 8).

Crucially, the morphology of the *D. toliapica* intraramal joint is strikingly similar to that of *Ichthyornis* and unlike that of extant neoavians bearing intraramal hinges. *D. toliapica* shows a ‘ball and socket’ articulation between the cotylar caudal end of the dentary/splenial and the rounded rostral condyle of the angular at the ventral edge of the joint, forming a synovial joint (Fig. 4B). A similar condylar synovial articulation is present *Ichthyornis*, in which the morphology of all these elements is remarkably reminiscent of the condition in *Dasornis* and contrasts with of the intraramal hinge of neoavians, in which the angular is mediolaterally thin rather than bulbous and inserts deep between the dentary and splenial (Fig. 2F; Zusi & Warheit 1992). A condylar joint, when present, is formed by a novel internal ossification situated dorsal to the angular, with its caudal condyle incorporated into the prearticular. The prearticular of *Dasornis*, as interpreted, inserts into a shallow groove between the dentary and splenial as in *Ichthyornis*, and is unlike the condition in extant neoavians with intraramal joints, in which the prearticular only contacts the splenial along its ventral edge (Fig. 2F).

#### Implications for pelagornithid phylogenetic relationships

Given the recent suggestion that several aspects of galloanseran cranial morphology that are shared with pelagornithids may be plesiomorphic for Neornithes (Mayr et al. 2018; Benito et al. 2022a), and the lack of key galloanseran synapomorphies in pelagornithids, the phylogenetic position of pelagornithids as total-group galloanserans must be questioned. Indeed, the possibility that pelagornithids fall outside of the avian crown group should not be dismissed, which would make them the only known lineage of stem birds to survive the end-Cretaceous mass extinction event (Longrich et al. 2011). Based on both well-established characters (including the intraramal joint and articular region as previously noted by Mayr and Rubilar-Rogers (2010) and Mayr et al. (2021)) and newly observed morphologies, the mandibles of pelagornithids also appear to be more similar to those of non-neornithine ornithurines than those of crown group birds. Further investigation of the arrangement of the lower jaw elements in pelagornithids with better preserved mandibles than *D. toliapica* is needed, as are further observations of extant neoavian birds with intraramal joints. This will be necessary to fully assess the hypotheses regarding pelagornithid mandible morphology presented here, and to determine whether similarities in mandible morphology between pelagornithids and non-neornithine ornithurines are convergent or symplesiomorphic.

### Inferring mandibular morphology of the ancestral neornithine

Despite the paucity of well-preserved crown bird skulls from the group’s early evolutionary history, a detailed understanding of crownward stem bird and crown bird morphology should allow us to draw a number of justified inferences about the ancestral condition of the neornithine mandible.

The fused mandibular symphysis of crown birds, a synostosis involving primary cartilage and chondroid bone (Bailleul et al. 2017), is considered an apomorphy of Neornithes (Cracraft et al. 2001; Turner et al. 2012). All known neornithines, with the possible exception of pelagornithids (Mayr & Rubilar-Rogers 2010), exhibit a fused mandibular symphysis, differing from the unfused condition of most non-neornithine ornithurines. The ancestral neornithine is inferred to have been toothless, with the edentulism characterising all extant birds predicted to have been the result of a single transition to toothlessness predating the origin of crown birds (Meredith et al. 2014; Wang et al. 2017; Louchart et al. 2021), although repeated acquisitions of avialan edentulism (Louchart & Viriot 2011; Wang et al. 2017; Brocklehurst & Field 2021) raise the possibility that palaeognath and neognath edentulism could have arisen independently. The absence of a predentary in Neornithes is potentially linked to the fusion of the mandibular symphysis and lack of teeth; a predentary is only known from toothed euornithines, and teeth combined with a toothless premaxilla and unfused mandible are thought to have been key to its kinetic function (Bailleul et al. 2019). A fused symphysis, edentulism, and the lack of predentary are therefore inferred to be synapomorphies diagnosing the avian crown group, Neornithes (Zhou & Zhang 2005; Turner et al. 2012).

Most extant neornithines, including palaeognaths, exhibit a tricondylar articulation of the quadrate with the mandible and thus three articular cotyles (lateral, medial, and caudal (Fig. 3D); Ericson 1997). A bicondylar articulation, with only two cotyles on the dorsal surface of the articular (lateral and medial), is present in all galloanserans (Weber & Hesse 1995; Ericson 1997; Mayr et al. 2018), *Asteriornis*, *Vegavis,* Pelagornithidae (Kuo et al. 2023) and some neoavian taxa such as Columbidae (Clarke et al. 2016), *Opisthocomus* (Ericson 1997) and the extinct gruiform *Aptornis* (Weber & Hesse 1995). It has previously been assumed that the ancestral state of the neornithine mandible was tri-cotylar (Thulborn 1984; Elzanowski 1995; Weber & Hesse 1995). Evidence for this, however, is not unambiguous, and it has similarly been suggested that two cotyles could be the plesiomorphic condition of Neornithes (Ericson 1996; Mayr et al. 2018), with some phylogenetic analyses suggesting independent transitions to a tricondylar articulation in Palaeognathae and Neoaves (Field et al. 2020). This hypothesis is supported by the presence of only two cotyles in crownward stem-birds such as *Ichthyornis* (Clarke 2004; Field et al. 2018), although three cotyles are present in the mandibles of hesperornithine birds (Bell & Chiappe 2020), often recovered in a phylogenetic position crownward to Ichthyornithes (e.g. Bell & Chiappe 2016; Benito et al 2022a,b). Additionally, recent analyses suggest that aspects of the palatal morphology of Galloanserae may retain the plesiomorphic condition of Neornithes (Benito et al. 2022a; Field et al. 2025; Benito et al. 2025); this may also hold true regarding the morphology of the quadrate-articular joint. Benito et al. (2022a) show that several traits of the palaeognath palate long assumed to represent the plesiomorphic neornithine condition are instead derived and highly variable; the existence of a high level of variation in the shape and prominence of the three articular cotyles of palaeognaths could be indicative of a similar scenario regarding the quadrate-mandible articulation (Elzanowski 1977; Houde & Haubold 1987; Bertelli et al. 2014). Furthermore, the tricondylar articulation of palaeognaths and Neoaves are noticeably different (Fig. 4B), as is the remarkably divergent morphology of the mandibular condyles of their quadrates (Kuo et al. 2023, 2024), consistent with the interpretation that a tricondylar articulation evolved convergently in both clades. Following recent insights (Mayr et al. 2018; Field et al. 2020; Benito et al. 2022a; Kuo et al. 2023), we suggest that two mandibular cotyles (and a bicondylar quadrate) likely represents the ancestral condition for Neornithes.

Large and well-developed retroarticular processes are present in Galloanserae and some neoavian taxa including flamingos and some members of Charadriiformes, Gruiformes, and Telluraves (Beecher 1951; Ericson 1997; Livezey 1997; Mayr 2005; Mayr 2022), although retroarticular morphology in Neoaves has generally been considered distinct from that of galloanserans (Livezey 1997). The absence of a large retroarticular has been considered the ancestral condition for Neornithes (Weber & Hesse 1995), which is supported by its absence in crownward stem-birds such as *Ichthyornis* and Hesperornithes (Clarke 2004; Martin & Naples 2008).

In addition to these well-documented morphologies, we found several new features which we suggest may have characterised the ancestral neornithine mandible. A cluster of morphological characteristics appear common to anseriforms, pelagornithids and, in some instances, *Vegavis.* These include an unforked dentary, a straight (not ventrally concave) mandibular profile, and a rostrally extensive prearticular (although the presence of this feature in *Vegavis* is unknown). A contrasting suite of characters is found in galliforms, palaeognaths and *Asteriornis*, including variably caudally forked dentaries, comparatively small splenial and prearticular bones, and a straight dorsal margin of the mandible lacking a prominent mandibular apex. Further analyses are required to assess which of these morphologies, or combinations of these characters, represent the ancestral neornithine condition; however, given that the first suite of characters is also found in crownward stem-birds including both *Ichthyornis* and Hesperornithes (Martin & Naples 2008), we suggest that these are most likely to be ancestral for Neornithes, with the second suite of characters shared by galliforms and palaeognaths being independently derived.

## Conclusions

Our data raise new questions regarding the affinities of several phylogenetically controversial fossil birds and emphasise the importance of detailed comparative investigations of morphological complexes such as the mandible for revealing new insights into the early evolutionary history of Neornithes. We find that mandibular morphology supports galloanseran and not palaeognath affinities for *Asteriornis* and, conversely, raises doubts about the galloanseran and neornithine affinities of *Vegavis* and pelagornithids. Future investigations on a broader phylogenetic scale, incorporating these anatomical observations and a wider range of fossil birds into quantitative phylogenetic analyses, should continue to help clarify the phylogenetic affinities of controversial fossils near the origin of the avian crown group.

## Declarations

### Ethics approval and consent to participate

No permissions were needed for collection of specimens during this study- all specimens used were already accessioned museum specimens and data collected with permission from the relevant institutions.

### Consent for publication

Not applicable

### Availability of data and materials

The CT datasets generated during the current study are available from the corresponding author on reasonable request.

### Authors’ contributions

A.H.C.: Conceptualization, Investigation, Methodology, Visualization, Writing– Original Draft Preparation. J.B.: Investigation, Visualization, Writing – Review & Editing. A.C.: Writing – Review & Editing. D.T.K.: Writing – Review & Editing. D.J.F.: Conceptualization, Supervision, Writing – Review & Editing.

## Acknowledgements

We thank Matthew Lowe (UMZC), Keturah Smithson (Cambridge Biotomography Centre), Mario Urbina and Raffo Varas-Malca (Universidad Nacional Mayor de San Marcos), and Mike Day (NHM) for assistance with specimen access and imaging.

