## Supplementary Material for "Mandibular morphology clarifies phylogenetic relationships near the origin of crown birds"

Supplementary Materials

| **Palaeognathae** | **Specimen Number** |
| --- | --- |
| *Struthio camelus* | UMZC uncatalogued |
| *Dromaius novaehollandiae* | UMZC uncatalogued |
| *Tinamus solitarius* | UMZC uncatalogued |
| **Anseriformes** |  |
| *Chauna charavia* | UMZC 12.Anh.2.a.3 |
| *Thalassornis leuconotus* | UMZC uncatalogued |
| *Anser fabalis* | UMZC 12/Ana/4/e/14 |
| **Galliformes** |  |
| *Megapodius pritchardii* | UMZC 14/Meg/b/g/2 |
| *Gallus gallus* | UMZC uncatalogued |
| **Neoaves** |  |
| *Morus bassanus* (juvenile) | UMZC 10.Sul.1.a.7 |
| *Diomedea exulans* | UMZC 9.Dio.1.h.8 |
| *Morus bassanus* (adult) | UMZC 262.C |

Supplementary Fig 1

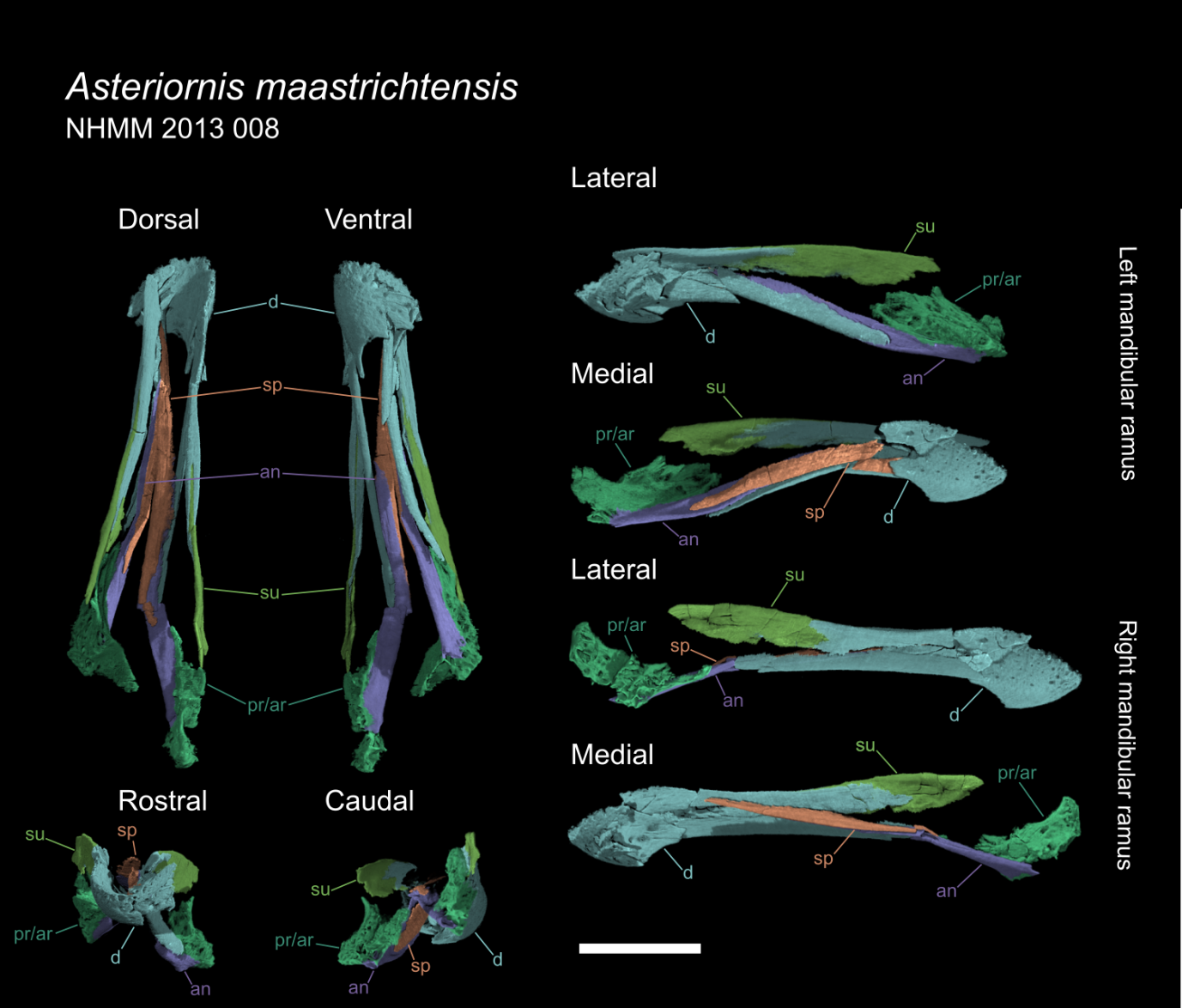

Supplementary Fig 2

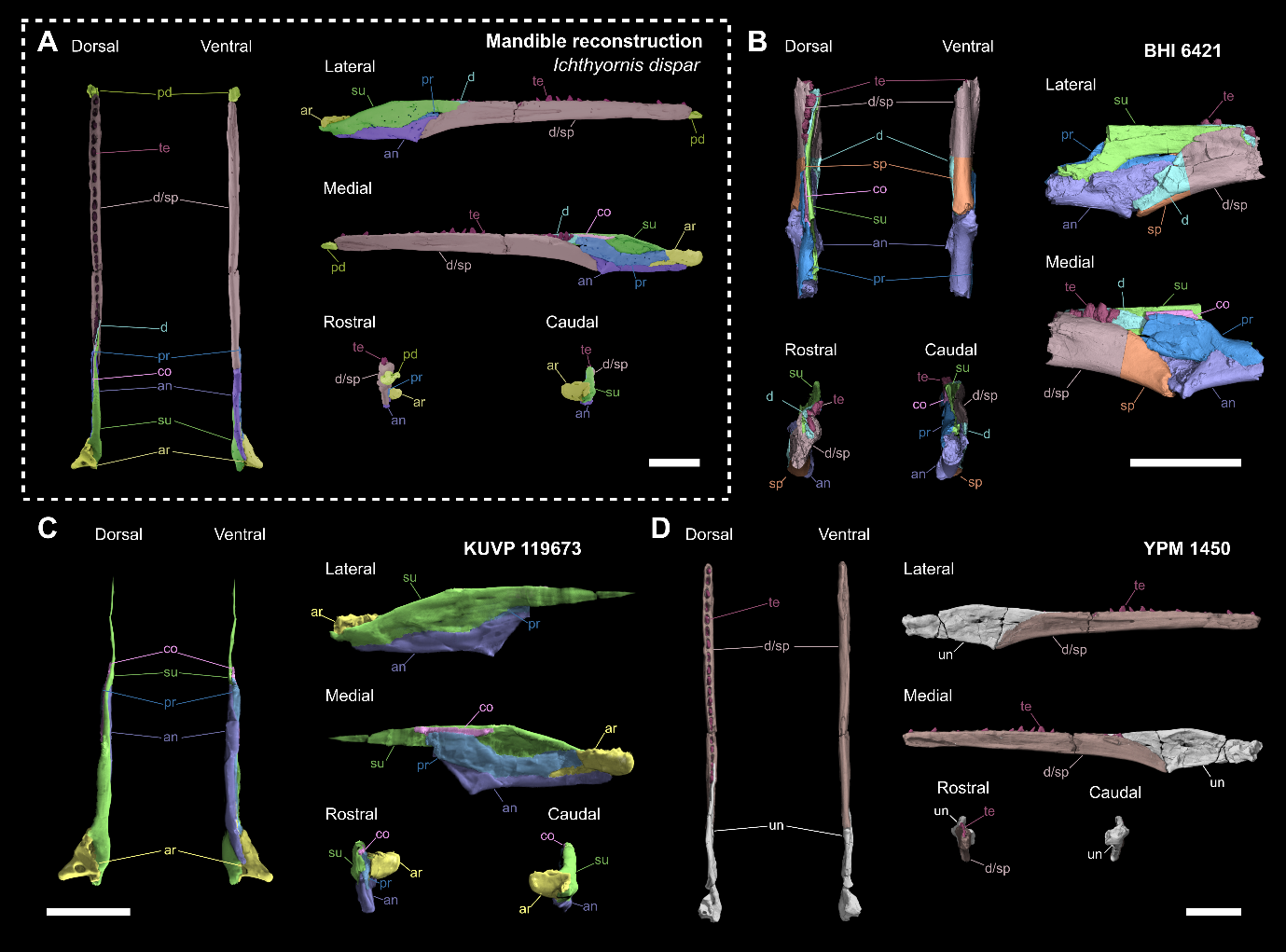

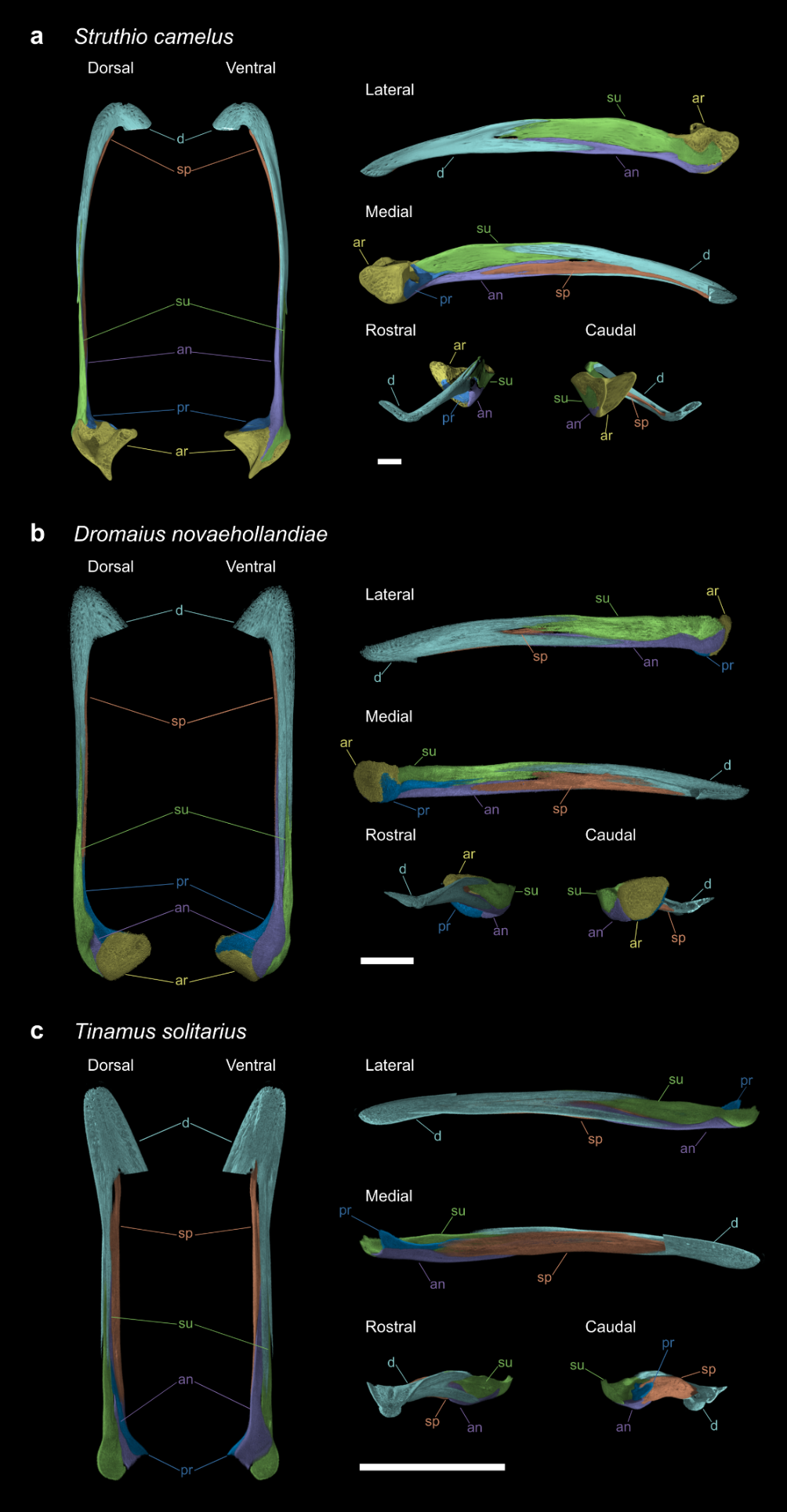
 Supplementary Fig 3

Supplementary Fig 4
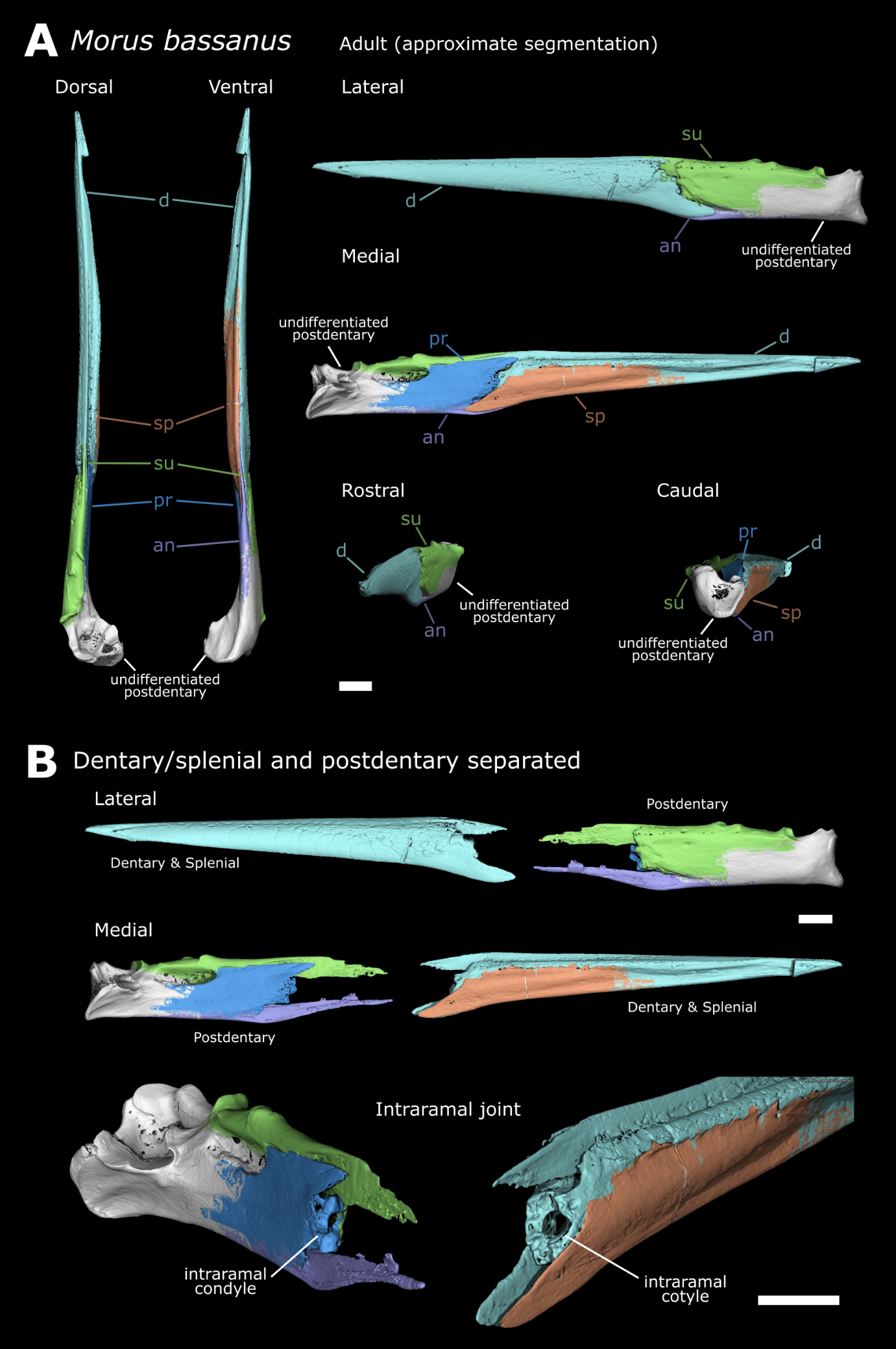

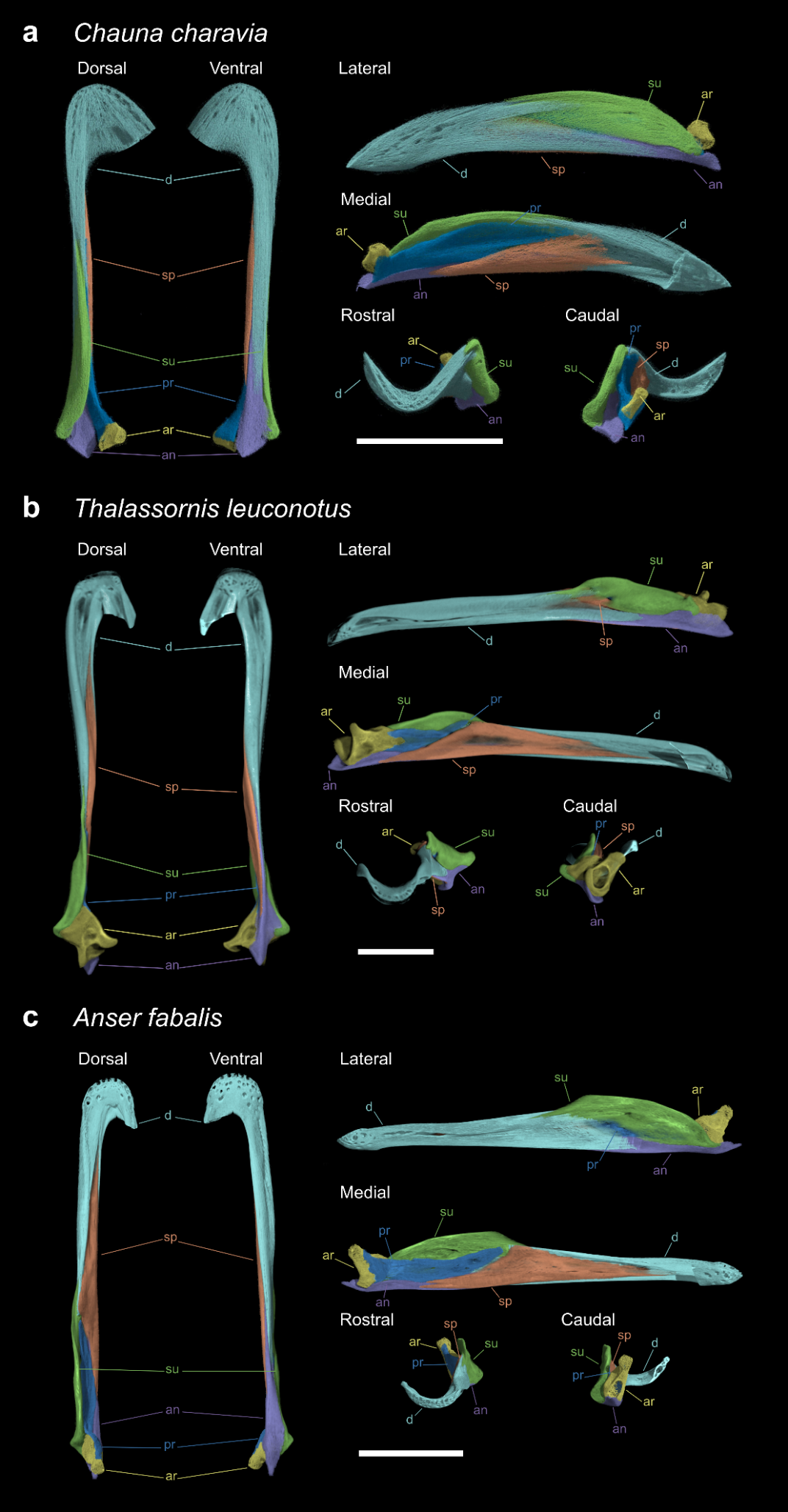
Supplementary Fig 5

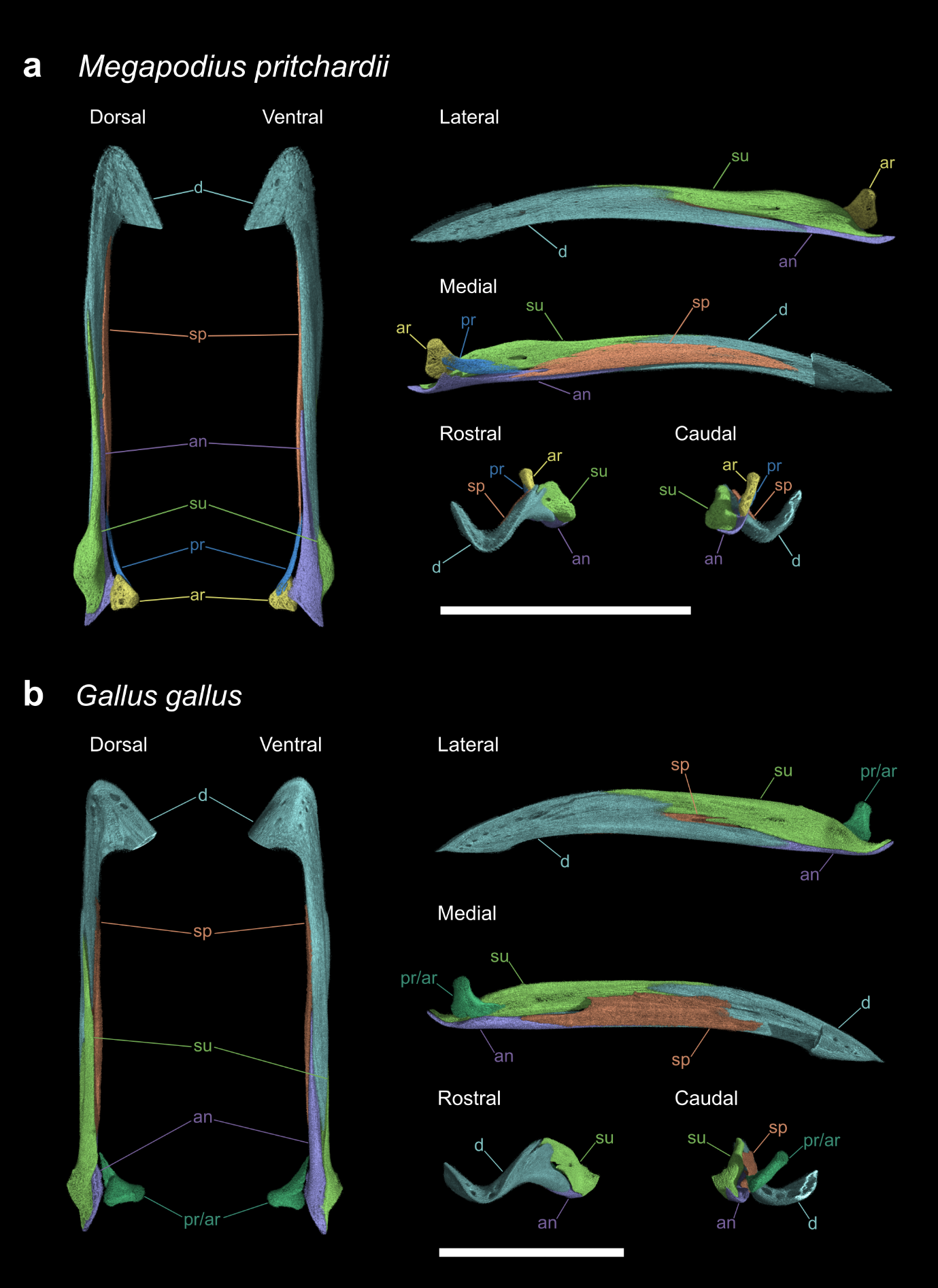
Supplementary Fig 6

Supplementary Fig 7

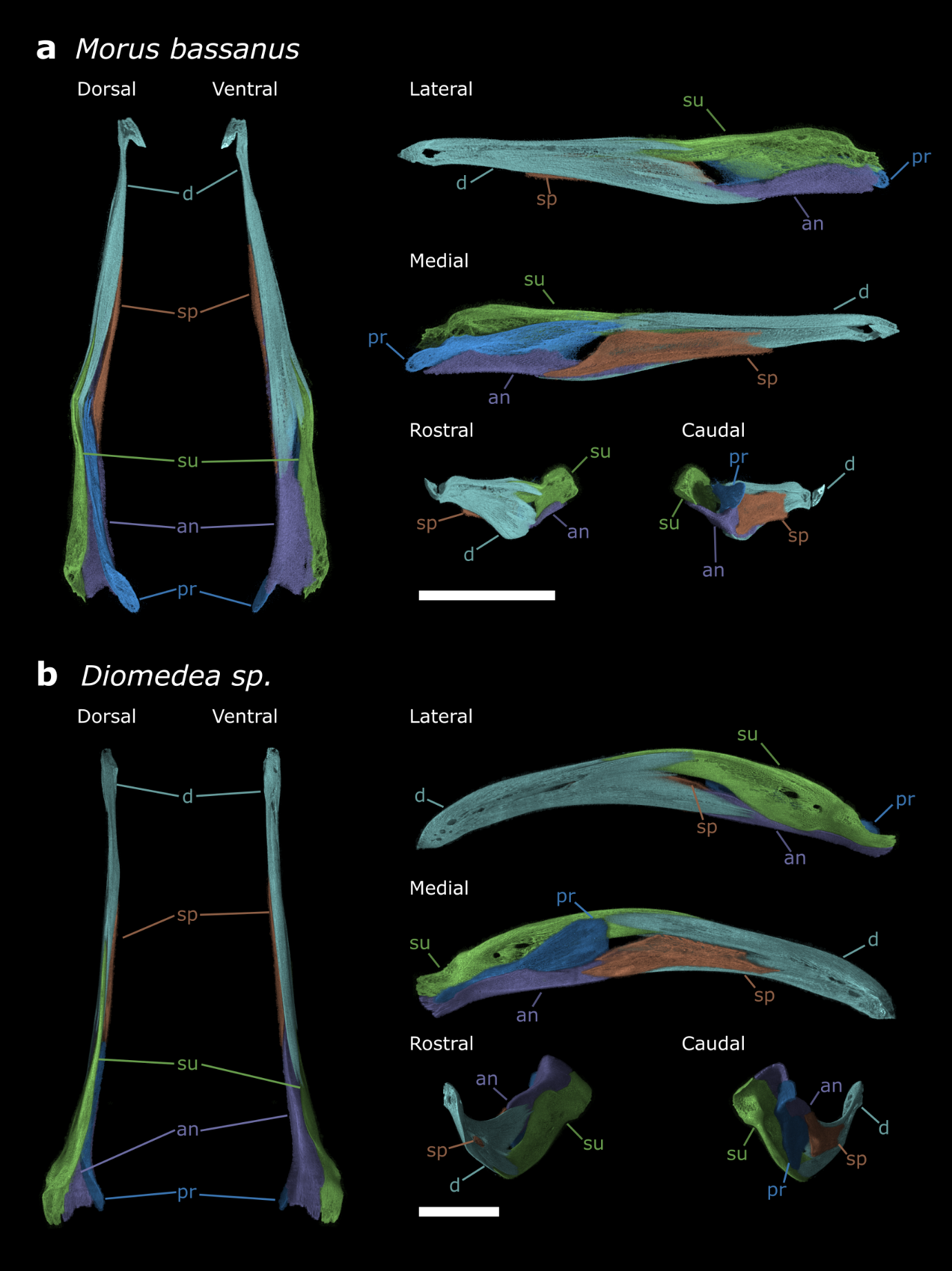

Supplementary Fig 8

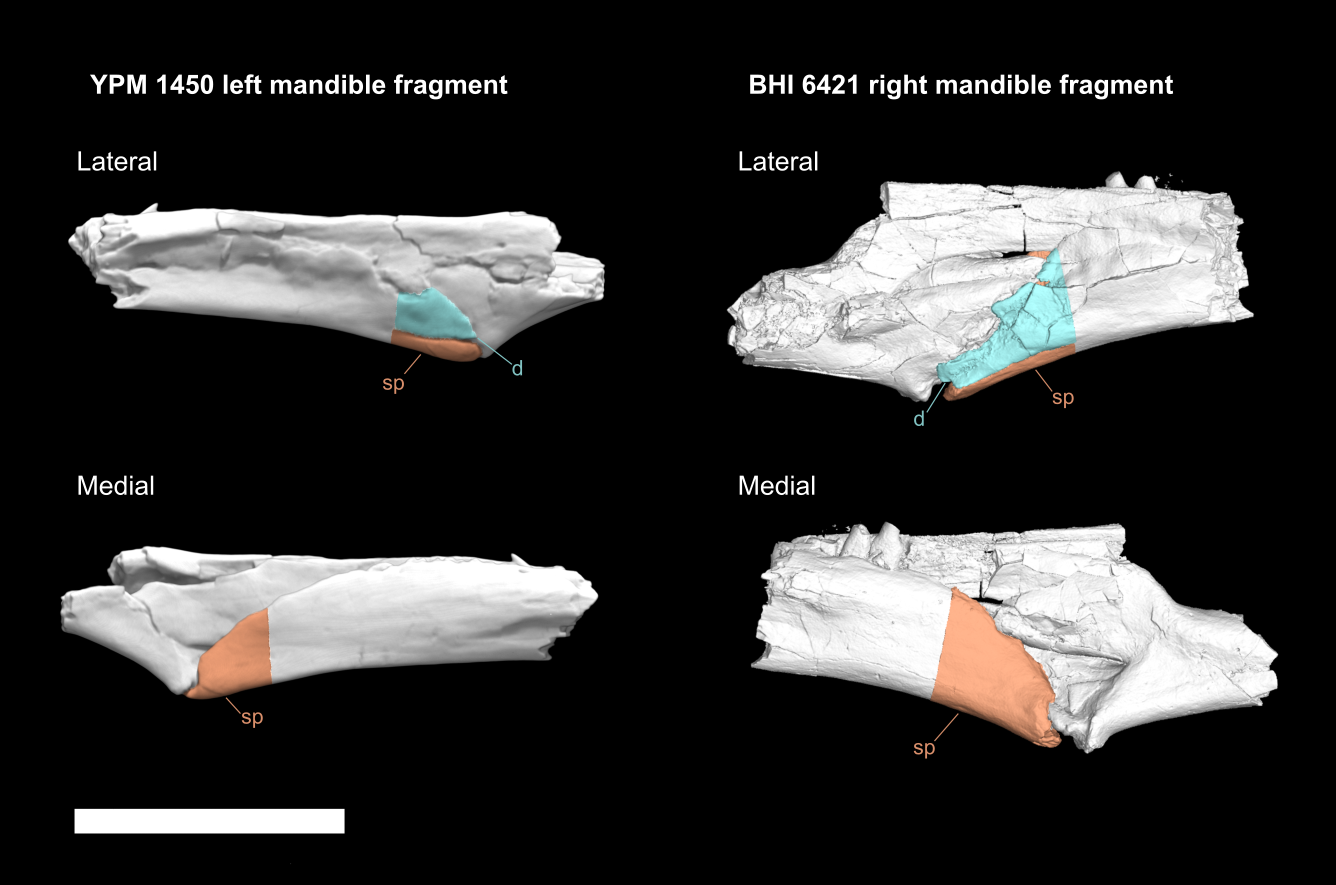

Supplementary Fig 9

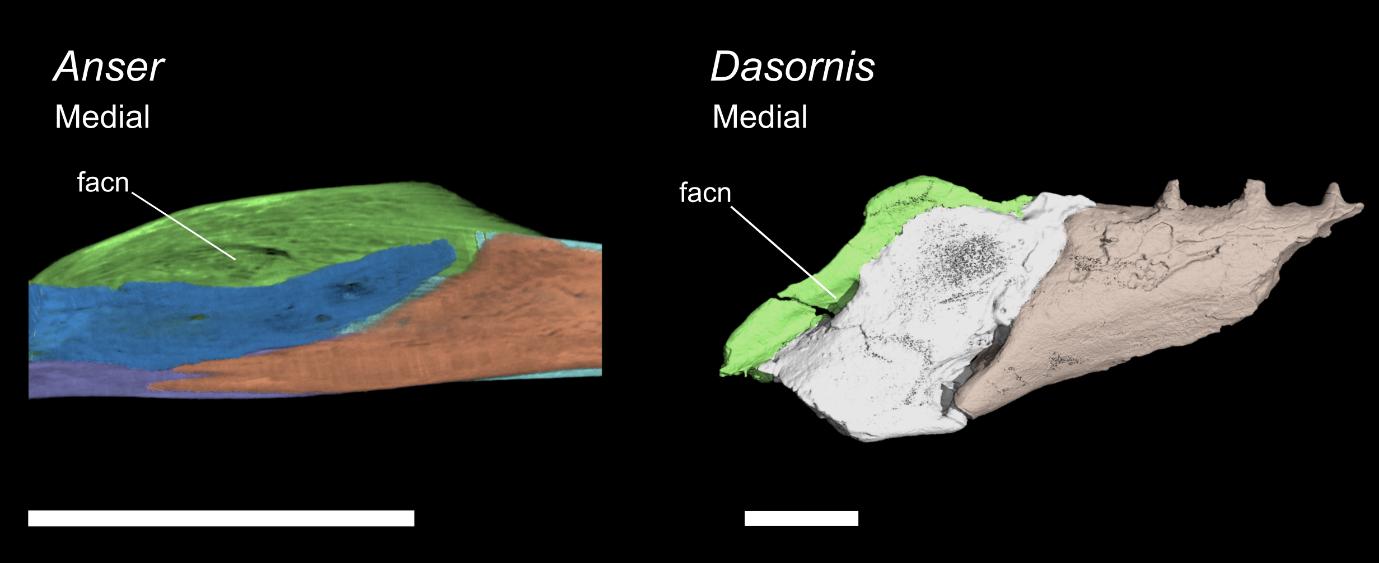

**Supplementary Table 1.** List of juvenile (and one adult) neornithine specimens CT scanned and used for mandibular anatomical comparisons.

**Supplementary Figure 1.** Mandible of *Asteriornis* holotype (NHMM 2013 008), as preserved. Whole mandible shown in dorsal, ventral, rostral and caudal views. Left and right rami shown separately in medial and lateral views. Prearticular and articular could not be distinguished, displayed as prearticular/articular combined. Scale bar equals 10mm.

**Supplementary Figure 2.** Right mandible of *Ichthyornis dispar* shown in dorsal, ventral, rostral, caudal, lateral and medial views. A) Reconstruction of mandible composed from multiple fossil specimens (BHI 6421, KUVP 119673, YPM 1450, AMNH FARB 32773), rescaled relative to BHI 6421. B) Segmentation of mandible section from *Ichthyornis* specimen BHI 6421. C) Segmentation of mandible section from *Ichthyornis* KUVP 119673. D) Segmentation of mandible section from *Ichthyornis* YPM 1450. Scale bars equal 10mm.

**Supplementary Figure 3.** Mandibles of juvenile palaeognaths shown in in dorsal, ventral, rostral, caudal, lateral and medial views. A) *Struthio camelus*. B) *Dromaius novaehollandiae*. C) *Tinamus solitarius-* this specimen is at an earlier ontogenetic stage and so does not exhibit an ossified articular bone. Scale bars equal 10mm.

**Supplementary Figure 4.** Mandible of adult *Morus bassanus* (Neoaves), approximately segmented. High levels of fusion in this adult specimen leads to uncertainty in parts of this segmentation; we have chosen to leave highly fused regions of the postdentary complex unsegmented. A) Whole ramus shown in dorsal, ventral, rostral, caudal, lateral and medial views. A) Mandible separated into dentary/splenial and postdentary complex, illustrating the anatomy of the intraramal joint. Ramus shown in in lateral, medial and oblique views. Scale bars equal 10mm.

**Supplementary Figure 5.** Mandibles of juvenile anseriforms shown in dorsal, ventral, rostral, caudal, lateral and medial views. A) *Chauna charavia*. B) *Thalassornis leuconotus*. C) *Anser fabalis*. Scale bars equal 10mm.

**Supplementary Figure 6.** Mandibles of juvenile galliforms shown in dorsal, ventral, rostral, caudal, lateral and medial views. A) *Megapodius pritchardii*. B) *Gallus gallus-* the prearticular and articular bones could not be distinguished in this specimen. Scale bars equal 10mm.

**Supplementary Figure 7.** Mandibles of juvenile neoavians with intraramal hinges shown in dorsal, ventral, rostral, caudal, lateral and medial views. A) *Morrus bassanus*. B) *Diomedea sp.* Both of these specimens are at an early ontogenetic stage and do not exhibit an ossified articular bone Scale bars equal 10mm.

**Supplementary Figure 8.** Mandibular fragments of Ichthyornis (YPM 1450 left mandible, BHI 6421) preserving an open suture between the dentary and splenial. The open suture between the dentary and splenial at their caudal ends is described in YPM 1450 left mandible by Clarke (2005) but not figured. This same suture us observable in part of BHI 6421. Scale bar equals 10mm.

**Supplementary Figure 9.** Partial mandible of *Dasornis* and similar mandibular portion of *Anser.* Part of left ramus in medial view, illustrating the fossa adius canalis neurovascularis in these taxa. Scale bar equals 10mm.
